# Precision Glycoform Engineering: Combining plant and *in vitro* systems for tailored biopharmaceutical production

**DOI:** 10.1101/2025.10.01.679750

**Authors:** Mijke R. Sweers, Ruud H.P. Wilbers, Arjen Schots, Pieter Nibbering

## Abstract

Protein biopharmaceuticals play a key role in providing effective, targeted, and personalized therapies for diverse diseases, while also preventing and mitigating a broad range of infections. N-glycosylation is a key post-translational modification influencing the biological activity of many protein-based therapeutics, yet structure–function relationships of N-glycans remain poorly understood due to challenges in producing homogeneous glycoforms. Current go-to production hosts, mammalian and yeast cells, often yield heterogeneous glycan profiles and require extensive genetic manipulation. Alternative production hosts such as the plant *Nicotiana benthamiana*, provide more homogeneous glycosylation and flexibility through transient expression, but are limited in the generation of certain glycoforms. *In vitro* glycoengineering can overcome these limitations but is time consuming and requires expensive resources. In this study, we show that by combining *in planta* and *in vitro* glycoengineering strategies, we can quickly produce a wide range of homogeneous glycoforms of pharmaceutical proteins with high mannose, paucimannose, hybrid and complex N-glycan structures. Using *N. benthamiana* as a transient expression host, we produced two pharmaceutical glycoproteins — the monoclonal antibody rituximab and the helminth vaccine candidate OoASP-1 — and modified them *in vitro* using *Escherichia coli* produced glycoenzymes. The combination of these two glycoengineering systems minimizes the amount of time and resources required, while maintaining high glycan homogeneity. This scalable, flexible, and cost-effective platform opens the door to glycan structure–function relationship studies and can support rational design of next-generation biopharmaceuticals.

## Introduction

As the global population approaches 10 billion by 2050 [1], the world will face a need for improved healthcare and more effective treatments to ensure sustainable living for humans and animals. This highlights the need for affordable and widely available biopharmaceuticals, like antibodies and vaccines, which can support both human and animal health through improved disease prevention and treatment. Most biopharmaceutical proteins receive post-translational modifications (PTMs), which are essential for the functionality and efficacy of most biopharmaceuticals. One of the most common PTMs is protein glycosylation, divided into O-glycosylation and N-glycosylation, of which the latter is most investigated in the context of biopharmaceuticals. N-glycosylation affects therapeutic performance in numerous ways. For monoclonal antibodies (mAbs), glycosylation is essential for activating effector functions through interactions with their Fc region [2,3]. N-glycosylation can also impact the elicitation of protective immunity by vaccines, as observed in the development of vaccines against HIV [4], SARS-CoV-2 [5] and the bovine nematode *Ostertagia ostertagi* [6]. Serum half-life of several biopharmaceuticals, such as erythropoietin [7] and follicle-stimulating hormone [8] is also affected by the presence or composition of N-glycans.

These studies highlight the increasing amount of knowledge we have about the functioning of glycans on biopharmaceuticals. However, as stated by the authors of the authoritative reference in the field of glycobiology, glycans are still considered ‘*the “dark matter” of the biological universe: a major and critical component that has yet to be fully incorporated into the “standard model” of biology*’ [9]. With the increase in analytical methods of glycans, the variety of glycans present on potential biopharmaceuticals, such as helminth glycoproteins [10], is being uncovered. However, the exact structure-function relationships of these glycans remain to be elucidated. By increasing our knowledge of glycan structure-function relationships we can exploit the potential of glycans for the rational design of therapeutic proteins, for instance by expanding the toolboxes for glycan-induced immunofocusing (as applied for HIV, influenza and SARS-CoV-2 [11]), immunotargeting (as applied in cancer immunotherapy [12]) or immunomodulation (as applied for treatment for inflammatory diseases [13]).

A major bottleneck for elucidating structure-function relationships of N-glycans is the ability to produce specific glycan variants (i.e., glycoforms). N-glycosylation is a complex, partially host specific process, which does not have a clear genetic footprint like a protein sequence encoded in DNA [14]. The process involves the co-translational attachment of N-glycans in the endoplasmic reticulum to an asparagine that is part of a N-glycosylation motif (N-X-S/T (X≠P)). The glycoprotein is then transported to the Golgi. From that point onwards, glycan processing pathways differ between species, and involve the action of multiple glycosidases and glycosyltransferases (glycoenzymes) that can trim or elongate the oligosaccharide chain [15]. The final N-glycan attached to a protein is the result of the complex interplay between, amongst others, the subcellular localization of the glycoenzymes in the Golgi, the availability of nucleotide sugar donors, the transport rate of the protein and the accessibility of the N-glycosylation motif(s) due to protein conformation [16–19].

To ensure product quality, current biopharmaceutical research has mainly focused on genetically engineering i.e., glycoengineering, commonly used expression hosts to have more ‘human-like’ N-glycans [20–23]. These glycoengineered production hosts have improved the production of many biopharmaceuticals and limit batch-to-batch variability [24]. Nevertheless, their inability to generate rare glycans limit their use in studies into the detailed structure-function relationship of N-glycans.

To facilitate such research, a precision glycoform engineering system is required than can generate a wide variety of homogeneous glycoforms [19,21,22][24]. Mammalian production systems struggle to fulfil this requirement due to N-glycan macro- and microheterogeneity. For example, CHO produced mAbs can be decorated with a mixture of over 50 different N-glycans [25], and N-glycans on HEK293 produced IL-22 are much more heterogeneous than on the plant-produced equivalent [26]. Though glycoengineered strains of mammalian cells yielding homogeneous glycans have been developed [22,23,27], their reliance on stable transformation makes them less adaptable to fluctuating research needs. Transient glycoengineering in plants (such as *Nicotiana benthamiana* [28]) is suitable in this respect, as it is cheap, fast, scalable and can generate relatively homogeneous N-glycans compared to mammalian production systems [29,30]. However, glycoengineering of *N. benthamiana* is limited in the generation of certain glycoforms due to plant specific glycosidases [31,32]. To circumvent limitations of *in vivo* glycoengineering, *in vitro* glycoengineering is often applied when specific and homogeneous glycans are desired.

*In vitro* glycoengineering is a form of downstream processing, where glycans are trimmed or elongated post-production [33]. As this is a tightly controllable process, it generally allows for the generation of more homogeneous glycans [34–36]. However, most studies focus solely on the introduction of galactose or sialic acid on complex glycans. Mannosylated glycans, though pharmaceutically relevant (as reviewed by Li et al. [37]) are not often generated *in vitro*. Additional drawbacks of *in vitro* glycoengineering include the effect of homogeneity of the *in vivo* produced starting material on final product homogeneity [38], and costs. *In vitro* glycoengineering is expensive compared to *in vivo* glycoengineering, as it requires production and purification of each individual glycoenzyme, and the addition of expensive nucleotide sugar donors [39,40]. There currently is no suitable production platform for the quick and resource efficient generation of homogeneously glycosylated biopharmaceuticals. The establishment of such a platform would facilitate studies into the structure-function relationships of glycans, and thereby enlarge the toolbox for glycan-induced biopharmaceutical optimization.

In this study, we show that by combining *in planta* and *in vitro* glycoengineering strategies, we can simplify the glycoengineering pathway and produce a wide range of homogeneous glycoforms of pharmaceutical proteins covering high mannose, paucimannose, hybrid and complex N-glycan structures (**Figure 1B**). All glycoforms were generated on two model biopharmaceuticals that rely on correct N-glycosylation to exhibit their function; monoclonal antibody rituximab [3] and helminth vaccine candidate *Ostertagia ostertagi* activation associated protein 1 (OoASP-1) [6]. Both proteins were produced efficiently in *N. benthamiana* using an *Agrobacterium*-mediated glycoengineering approach. The plant produced proteins were further modified *in vitro* with *Escherichia coli* produced glycoenzymes lacking their transmembrane region, to easily generate all biologically relevant glycoforms of rituximab and OoASP-1. The precision glycoform engineering system presented here enables the production of biopharmaceutical proteins with a variety of N-glycans and opens the door to investigating structure-function relationships to enhance biopharmaceutical performance.

**Figure 1.**
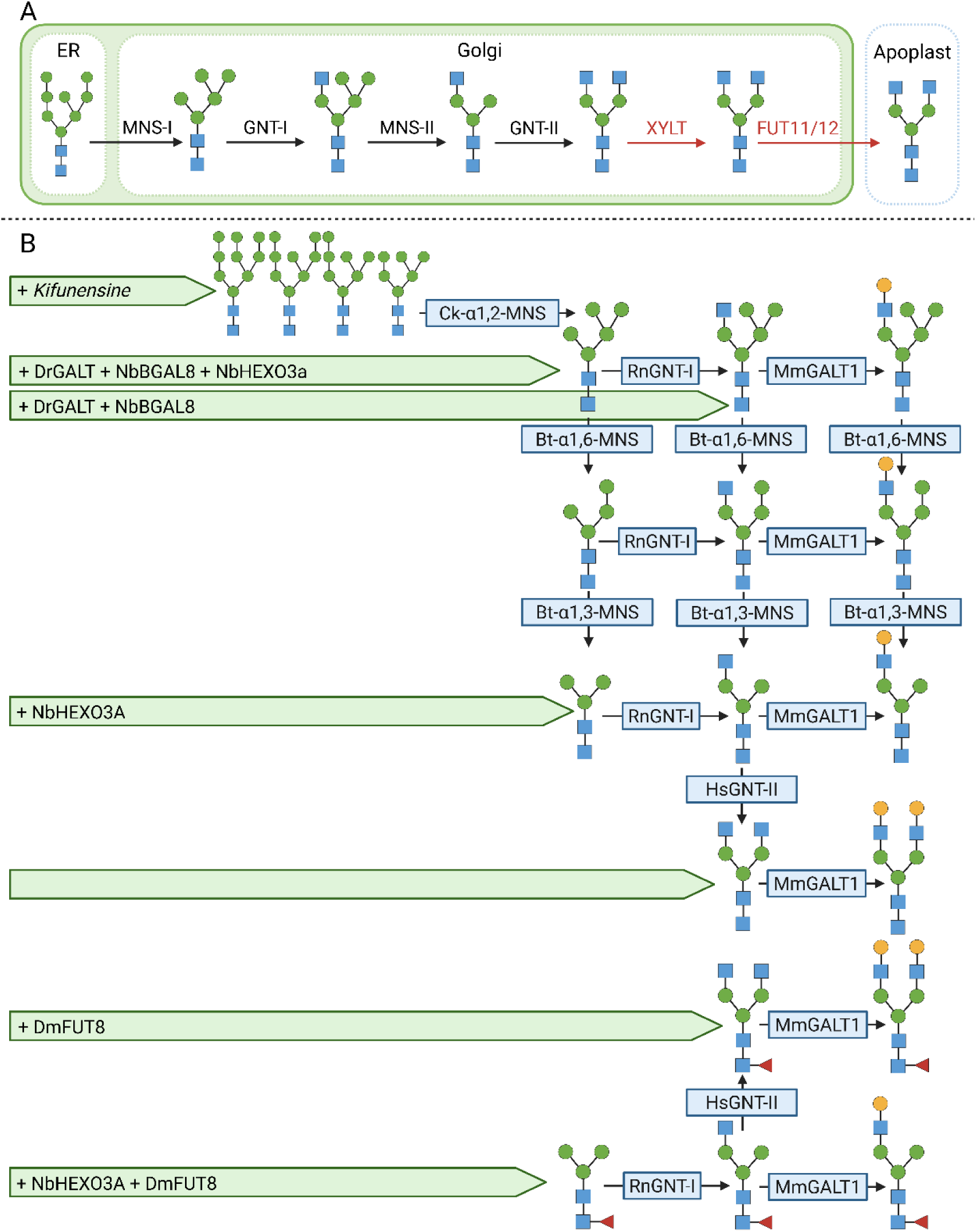
*In planta* and *in vitro* glycan processing pathways. **A** General glycan processing pathway of the *N. benthamiana* XT/FT plant line. **B** Glycan processing pathways than can be achieved through combining *in planta* and *in vitro* glycoengineering. Green arrows represent chemicals (*italics*) or glycoenzyme-encoding plasmids in *A. tumefaciens* that are infiltrated into *N. benthamiana* XT/FT plants to facilitate *in planta* glycoengineering. Arrows with blue blocks indicate *E.coli* produced enzymes that are used for *in vitro* glycoengineering reactions. Glycan structures are drawn according to symbol nomenclature for glycans (SNFG). Created in <u>BioRender</u>.

## Materials & Methods

### Generation of expression plasmids for OoASP-1, rituximab, and glycoenzymes

To facilitate *in planta* expression of rituximab, the variable regions of the heavy (γ) and light (κ) chains (γ = PDB: 8VGN_A, κ = PDB: 2OSL_B) (**Table S1**) were codon-optimized in-house [41] and synthetically produced by GeneArt, incorporating BspHI/NheI and NcoI/BsiWI overhangs, respectively. The variable regions were initially cloned into the shuttle vector pRAPa as previously described by Westerhof et al. [42], containing a dual Cauliflower Mosaic Virus (CaMV) 35S promoter, the signal peptide of the *Arabidopsis thaliana* chitinase gene (cSPN), and the constant region of either the gamma-1 heavy chain or kappa light chain. These constructs were then subcloned into the pHYG expression vector, as previously described by Westerhof et al. [42]. For expression of *Ostertagia ostertagi* activation associated protein-1 (OoASP-1), a pHYG vector containing the dual CaMV 35S promoter-driven cSPN-OoASP-1 expression cassette was used [6].

To generate expression plasmids for *E. coli* production of glycoenzymes, DNA sequences codon optimized for *E. coli* strain K12, lacking a signal peptide/transmembrane domain, were ordered according to **Table 1** and **Table S2** at GeneArt or integrated DNA technologies (IDT). The synthetic genes (**Table S2**) were cloned into the pCold DNA cold-shock expression system (Takara), upstream of the translation enhancing element (TEE), using the NEBuilder® HiFi DNA Assembly (Gibson) Cloning Kit and primers listed in **Table S3**. For Bt-α1,3-MNS and Bt-α1,6-MNS, the ordered linear synthetic fragments were directly assembled with a linear pCOLD fragment into a circular plasmid using the Gibson method.

**Table 1:** Enzymes produced in E. coli to facilitate the in vitro glycoengineering work.

| Enzyme name used here | Original enzyme name | Uniprot | Relevant literature | Expressed amino acids | N-terminal tags | Expression line |
| --- | --- | --- | --- | --- | --- | --- |
| Ck-α1,2-MNS | *Caulobacter* strain K31 GH47 (CkGH47) α1,2-mannosidase | NA | [43] | - | 6xHIS | BL21 |
| Bt-α1,3-MNS | *Bacteroides thetaiotaomicron* 1769 (Bt1769) α1,3-mannosidase | Q8A6V7 | [44,45] | Δ25-751 | 6xHIS | BL21 |
| Bt-α1,6-MNS | *Bacteroides thetaiotaomicron* 3994 (Bt3994) α1,6-mannosidase | Q8A0M7 | [44,45] | Δ25-743 | 6xHIS | BL21 |
| RnGNT-I | *Rattus norvegicus* Alpha-1,3-mannosyl-glycoprotein 2-beta-N-acetylglucosaminyltransferase | Q09325 | NA | Δ38-447 | Small Ubiquitin-like Modifier (SUMO)-6xHIS | BL21 |
| HsGNTI-II | *Homo sapiens* Alpha-1,6-mannosyl-glycoprotein 2-beta-N-acetylglucosaminyltransferase | Q10469 | NA | Δ31-447 | 6xHIS-Thioredoxin | Rosetta-Gami 2 |
| MmGALT1 | *Mus musculus* Beta-1,4-galactosyltransferase 1 | P15535 | NA | Δ44-399 | Maltose-Binding Protein (MBP)-TEV-6xHIS or SUMO-6xHis | BL21 |
| MmFUT8 | *Mus musculus* Alpha-(1,6)-fucosyltransferase | Q9WTS2 | NA | Δ107-575 | 6xHIS-NEXT | Rosetta-Gami 2 |

To facilitate *in planta* glycoengineering, several previously published glycosyltransferase and glycosidase expression cassettes were used. For incorporation of core α1,6-fucose, an expression plasmid for *Drosophila melanogaster* fucosyltransferase 8 (DmFUT8) was used [26]. For core α1,3-fucose, *Schistosoma mansoni* fucosyltransferase C (SmFucTC) was used [46]. To steer towards hybrid glycans, a β1,4-galactosyltransferase from *Danio rerio* with a CTS of rat α2,6-sialyltransferase under a 35S promoter (DrGALT) was used [47]. To cleave off *N*-acetylglucosamine and β1,4-galactoses, pHYG plasmids for expression of *N. benthamiana* hexosaminidase 3A (NbHEXO3A) [48] and *N. benthamiana* β-galactosidase 8 (NbBGAL8) [31] were used, respectively.

### Protein production and purification from *N. benthamiana*

To produce rituximab and OoASP-1 in *N. benthamiana*, the previously described expression plasmids were transformed to *Agrobacterium tumefaciens* strain MOG101. To enhance protein production, an *A. tumefaciens* strain harboring a construct for the expression of the viral silencing suppressor p19 [49] was used in each production. *Agrobacterium* culturing was performed as previously described by Wilbers et al. [47]. All cultures were resuspended in infiltration buffer (20 g/L sucrose, 5 g/L MS-salts, 1.95 g/L MES, pH 5.6, supplemented with 200 μM acetosyringone) to an OD600 of 0.5. To generate high mannose N-glycans, the infiltration suspension was supplemented with 0.5 µM kifunensine (Biosynth/Carbosynth) right before infiltration. Suspensions were infiltrated into the two youngest fully expanded leaves of five to six-week-old *N. benthamiana* plants with reduced expression of the XYLT and FUT11/12 (XT/FT [50]). After 5 days, leaves were harvested, and recombinant proteins were acquired via an apoplast fluid extraction as described by Wilbers et al. [26].

For purification of rituximab, apoplast fluids were dialyzed towards an equilibration buffer (20 mM PBS, 150 mM KCl, pH 7.0) using either SnakeSkin dialysis tubing (3500 MWCO, ThermoScientific) or Slide-A-Lyzer MINI Dialysis Units (3500 MWCO, ThermoScientific), depending on sample size. The samples were then allowed to bind to pre-equilibrated protein A resin (Abbexa) for 1 hour at 4°C while mixing, and washed 3 times with 5 bed volumes of cold equilibration buffer (20 mM PBS, 150 mM KCl, pH 7.0). Rituximab was eluted with 1 bed volume of 100 mM glycine (pH 3.0) into 1:50 bed volume of 1 M Tris-HCl (pH 9.0) to neutralize the buffer.

For purification of OoASP-1, apoplast fluids were dialyzed to cation exchange equilibration buffer (50 mM sodium acetate buffer, pH 4.4). The dialyzed OoASP-1 samples were allowed to bind to pre-equilibrated HS POROS® 50 strong cation exchange resin (ThermoFisher) for 30 minutes at 4°C and subsequently washed three times with equilibration buffer. The samples were eluted with equilibration buffer supplemented with 1 M NaCl.

The purified OoASP-1 and rituximab samples were dialyzed towards 50 mM HEPES pH 6.8 as previously described and were then used for subsequent enzymatic modifications.

### Glycoenzyme production in *E. coli*

The pCOLD plasmids harboring either the Ck-α1,2-MNS, Bt-α1,3-MNS, Bt-α1,6-MNS were transformed in chemical competent BL21 (Takara) cells and selected on appropriate selection media. A single colony was picked, and the cells were grown to an OD600 of 0.6-0.8 in LB medium at 37°C and cooled on ice for 30 minutes. Then, the gene expression was induced by adding IPTG to a final concentration of 0.5 mM and the cells were incubated at 15°C while shaking at 200 RPM for 16-24 hours. The cells were pelleted and resuspended in ice-cold extraction buffer (25 mM HEPES, 300 mM NaCl, 25 mM imidazole, pH 7.4). The cells were then bead-milled with glass beads (500-750 µm) for 8 minutes at 30 m/s. Subsequently, the broken cells were pelleted by spinning them down at 18.000g for 5 min at 4°C, and the supernatant was added to pre-equilibrated HisPur™ Ni-NTA Resin and incubated for 30 min at 4°C while mixing. The resin was washed 3 times with 4 volumes of ice-cold wash buffer (25 mM HEPES, 300 mM NaCl, 25 mM imidazole, pH 7.4) and then the enzymes were eluted with 1 bed volume of elution buffer (25 mM HEPES, 300 mM NaCl, 250 mM imidazole, pH 7.4). RnGNT-I and MmB4GALT were produced and purified similarly, except that after cell disruption, 3-((3-cholamidopropyl) dimethylammonio)-1-propanesulfonate (CHAPS) was added to 1% (w/v) and the lysate was mixed for 1 h at 4 °C to enhance enzyme solubility.

The pCOLD plasmids harboring the HsGNT-II and MmFUT8 genes were transformed into electrocompetent Rosetta-Gami 2 (DE3) pLysS (Novagen) and selected on appropriate selection media. The enzymes were produced as described in the previous paragraph, except induction with IPTG was prolonged to 48 hours. MmFUT8 was purified according to the previously described paragraph without CHAPS, while HsGNT-II was purified including CHAPS incubation.

Finally, the enzymes were dialyzed towards 25 mM HEPES, 100 mM NaCl pH 6.8 and used for the subsequent enzymatic reactions.

### Coomassie blue staining and western blot

All protein purifications were visualized via Sodium Dodecyl Sulfate Polyacrylamide Gel Electrophoresis (SDS-PAGE) followed by Coomassie blue staining. SDS-PAGE was performed using 12% (w/v) Bis-Tris mPAGE gels (MerckMillipore) in 1x MES buffer (50 mM MES; 50 mM Tris-base; 3,465 mM SDS; 1,025 mM Na-EDTA) under reducing conditions. Coomassie staining was performed as previously described by van der Kaaij et al. [31].

The purification of MmFUT8 and HsGNT-II was visualized via an anti-His tag western blot. Proteins were separated via SDS-PAGE as previously described and gels were transferred to a 0.45 μM PVDF membrane using a Turbo transfer blotter (BioRad) at 25 V, 1.3 A for 15 min. PVDF membranes were blocked for 1 hour at room temperature with 3% (w/v) bovine serum albumin (BSA) in 1x Phosphate-Buffered Saline with 0.1% Tween 20 (PBST). The PVDF membranes were then blotted with an HRP-conjugated anti-His antibody (Miltenyi Biotec, 130-092-785) with a 1:5000 dilution dissolved in 1.5% (w/v) BSA in PBST.

### *In vitro* glycoengineering reactions

The *in vitro* glycoengineering reactions were performed in a sequential manner as described in **Figure 1B**. All the enzymatic reactions were performed in 50 mM HEPES, 1 mM CaCl_2,_ pH 6.8. For MmGNT-I, HsGNT-II and MmGALT1, 10 mM MnCl_2_ was added as co-factor. For RnGNT-I and HsGNT-II UDP-GlcNAc (Merck) was used as sugar donor while the MmGALT1 reactions contained UDP-galactose (Merck) as sugar donor in the amounts described in **Table 2**. MmFUT8 reactions contained GDP-fucose (Biosynth) as sugar donor. Low purification yields prevented measurement of MmFUT8 concentrations and calculation of enzyme-to-substrate ratios. As a result, purified and dialyzed MmFUT8 comprised 50% of the reaction volume.

**Table 2:**
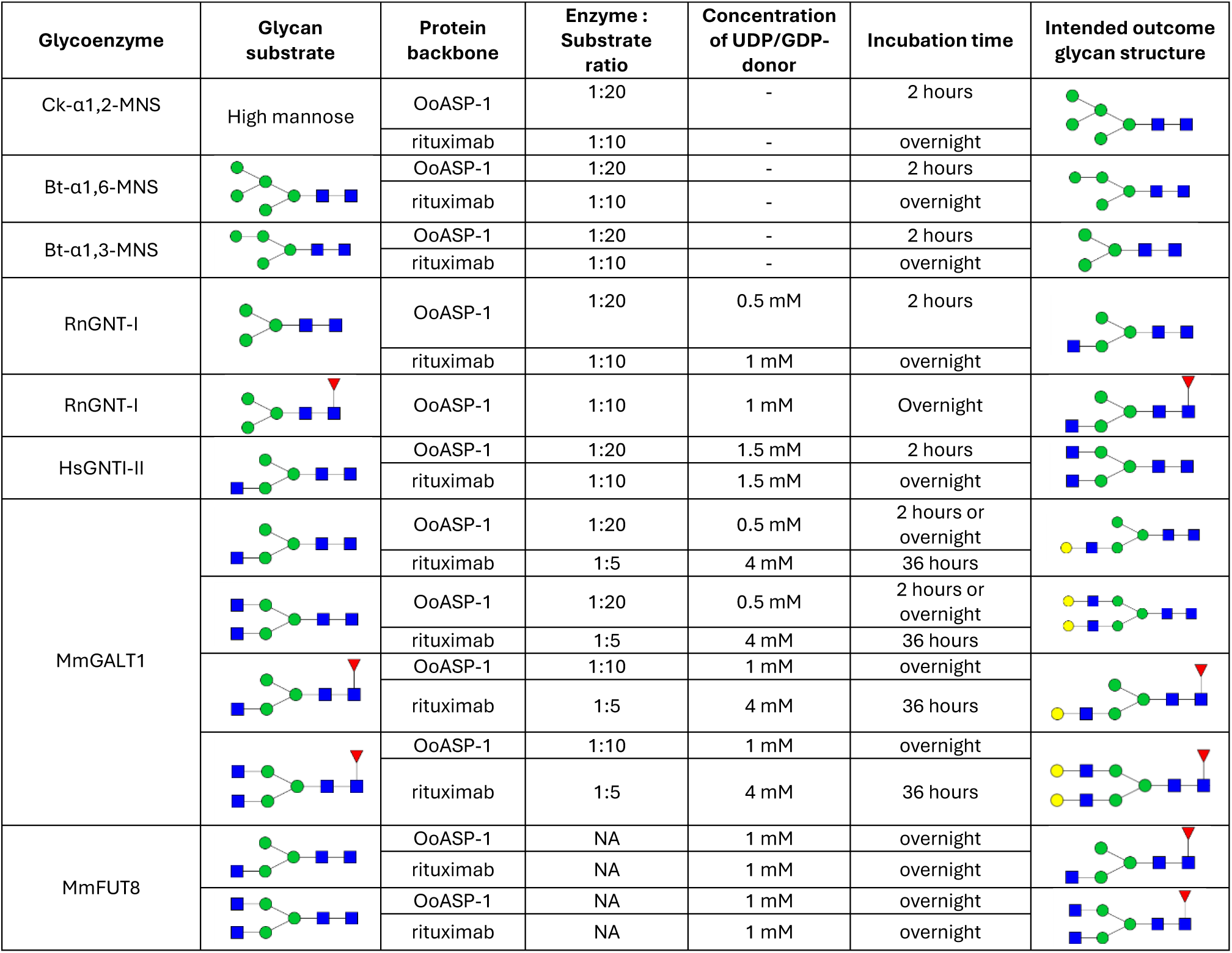
*In vitro* enzymatic conditions for glycan modification of OoASP-1 and rituximab.

### *In planta* glycoengineering

When *in planta* glycoengineering was needed, kifunensine was omitted from the infiltration buffer during protein production. Instead, *A. tumefaciens* strains carrying relevant glycoenzyme expression plasmids (green bars of **Figure 1B**) were combined in the infiltration suspensions for OoASP1 and rituximab. All glycoengineering constructs were resuspended to an OD600 of 0.5, except for DmFUT8, which was resuspended to an OD of 0.1. The suspensions were infiltrated into five-to six-week-old XT/FT plants. All other procedures regarding agroinfiltration, harvesting and purification remained as previously described.

### MALDI-TOF sample preparation & measurement

To analyze the N-glycan composition of the glycoengineering proteins, 50 µg of OoASP-1 or 100 µg of rituximab was deglycosylated according to the PNGase F protocol of New England Biolabs (NEB, #P0704). Released N-glycans were purified using BAKERBOND^TM^ Octadecyl SPE columns (J.T Baker) and Supelclean^TM^ ENVI-Carb SPE columns (MERCK) according to the suppliers’ protocols. Carbon SPE eluates were dissolved in 5 μM NaCl, of which 1 μl was spotted onto a ground steel MALDI target plate (MTP 384, Bruker Daltonics) with 1 μl of matrix solution (20 mg/ml 2,5-dihydroxybenzoic acid (Bruker Daltonics) in 30% ACN) and dried under a stream of warm air. Dried spots were analyzed using an AutoFlex Max MALDI/TOF MS system (Bruker Daltonics) operated by FlexControl software. For calibration, 1 mg/ml of maltodextrin (in H_2_O) was used. Samples were measured in positive mode using the reflector, with a delayed extraction time of 140ns and acceleration voltage of 19 kV over a range of 700-3500 Da, with deflection up to 650 Da. The molecular masses were determined using FlexAnalysis (Bruker Daltonics) without smoothing or baseline correction. Spectra were cropped and annotated using mMass [51] and GlycoWorkBench [52].

## Results

To establish a versatile and cost-effective *in vitro* glycoengineering platform, we aimed to produce a broad range of glycoenzymes in *Escherichia coli*, capable of modifying high-mannose N-glycan structures into more complex N-glycans on our target proteins. We chose high mannose N-glycans as a starting point for *in vitro* engineering as high mannose N-glycans represent the initial glycan structures entering the Golgi apparatus, and thus can be used to test an *in vitro* glycosylation system that mimics the eukaryotic glycosylation pathway. As protein substrates we chose pharmaceutical proteins OoASP-1 and rituximab since both rely on hybrid or complex N-glycans for their functionality [3,6], thereby making them relevant targets for precision glycoform engineering. Both targets were produced in *N. benthamiana* XT/FT plants (**Figure 1A**), as this production host yields relatively homogeneous glycans compared to other eukaryotic hosts [26,29].

### *In planta* production of high mannose OoASP-1 and rituximab

High mannose OoASP-1 and rituximab were produced in plants by using the endoplasmic reticulum (ER)/Golgi apparatus mannosidase inhibitor kifunensine. This inhibitor is a potent and selective inhibitor of class I α-mannosidases that are involved in N-glycan processing and has been utilized across a diverse array of organisms and cell systems to produce high mannose N-glycosylated target proteins [53–55]. Both OoASP-1 and rituximab were extracted via apoplast extraction and subsequently purified using cation exchange and protein A chromatography, respectively (**Figure S1**). MALDI-TOF MS analysis confirmed the synthesis of a range of high-mannose N-glycans on both OoASP-1 and rituximab (**Figure 2A**). Rituximab predominantly featured Man_9_GlcNAc₂ to Man_7_GlcNAc₂ structures, whereas OoASP-1 exhibited a broader spectrum, extending down to Man_5_GlcNAc₂. Interestingly, both contained only minimal levels of processed N-glycans with single or double antennary GlcNAc, indicating near-complete inhibition of the plant native class I α-mannosidases under these conditions. The high mannose substrates were further used to test the *in vitro* glycoengineering platform.

**Figure 2.**
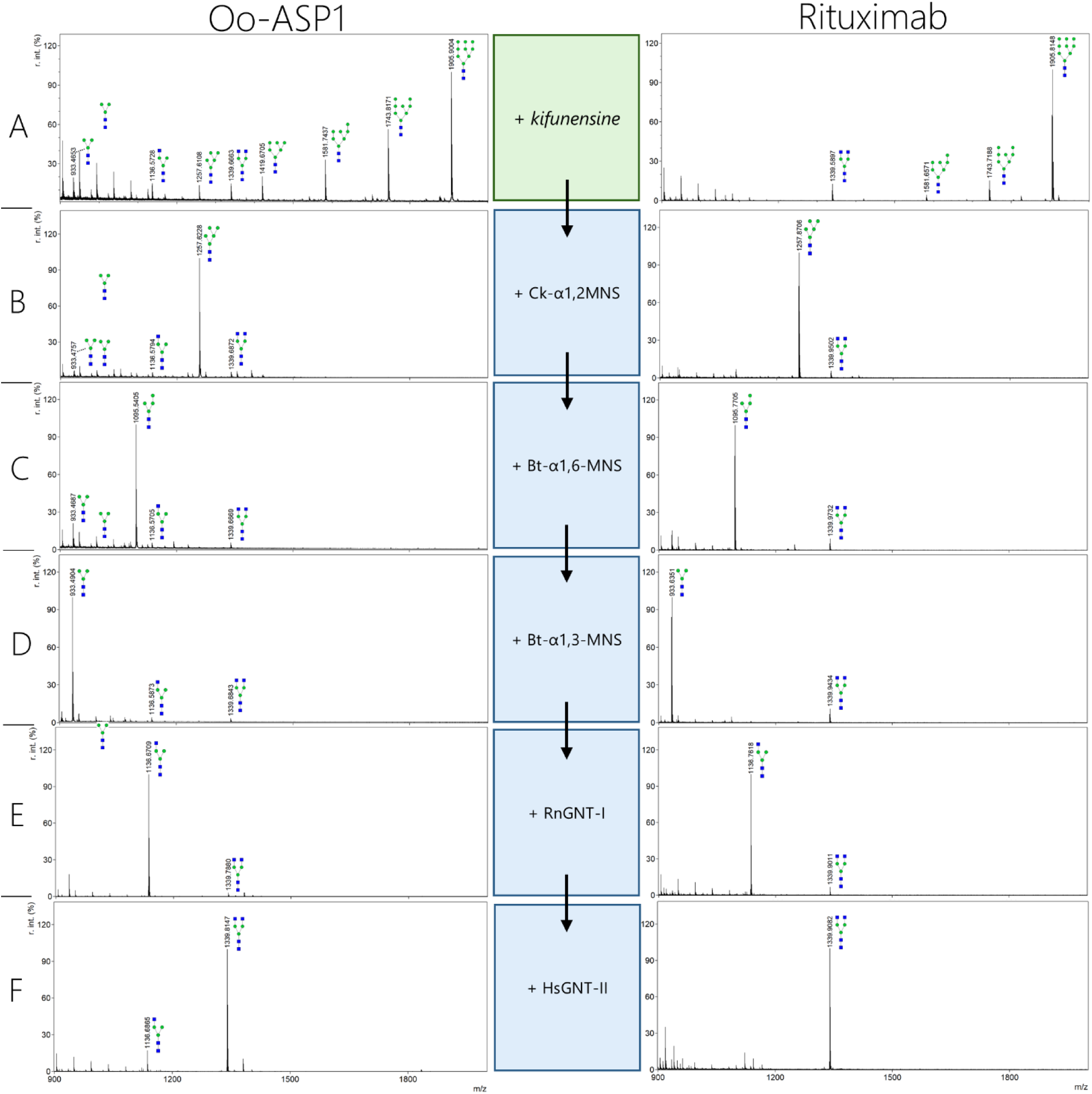
MALDI-TOF MS spectra of PNGase F released glycans generated on OoASP-1 and Rituximab through sequential *in vitro* glycoengineering. Plant-produced high-mannose OoASP-1 and Rituximab (Green box, **A**) were sequentially modified to reach a biantennary GlcNAc using *E. coli*–expressed glycoenzymes (Blue boxes, **B-F**). Black arrows indicate sequential *in vitro* reactions. Peaks that could not be labeled to glycans are left unannotated. Glycan structures are drawn according to symbol nomenclature for glycans (SNFG).

### *In vitro* glycoengineering of high mannose OoASP-1 and rituximab to complex N-glycans

Several glycoenzymes (Table 1) were produced in *E. coli*, providing a cost-effective platform for high-yield enzyme production while enabling essential post-translational modifications such as disulfide bond formation in engineered *E. coli* strains [56]. We chose the pCold DNA cold-shock expression system from Takara because it enables high-purity protein expression in *E. coli* at low temperatures, which reduces host protein expression, decreases protease activity, and helps prevent inclusion body formation [57]. All the enzymes described in **Table 1** were purifiable from *E. coli* (**Figure S2, S3 & S4**). However, the mammalian glycoenzymes showed no or low solubility under standard conditions. Solubility enhancing tags and the zwitterionic detergent CHAPS drastically improved the solubility of RnGNT-I, HsGNT-II and MmGALT1 enzymes (**Figure S2**). Unfortunately, the NEXT-tagged MmFUT8 showed low solubility and was only purifiable to low levels (**Figure S4**).

The *E. coli* produced glycoenzymes were then used to sequentially modify the plant-produced OoASP-1 and rituximab with high mannose N-glycans to generate different glycoforms. To reach a Man_5_GlcNAc₂ structure, the previously characterized Caulobacter α1,2-mannosidase (Ck-α1,2-MNS) was used [43]. The enzyme efficiently trimmed N-glycans on both OoASP-1 and rituximab to Man_5_GlcNAc₂ (**Figure 2B**). However, rituximab required higher amounts of enzyme and longer incubation times to achieve the desired glycoform compared to OoASP-1 (**Table 2 & Figure S5**). Similar observations were made for subsequent trimming to Man_4_GlcNAc₂ and Man_3_GlcNAc₂ on rituximab by α1,6-mannosidase (Bt-α1,6-MNS) and α1,3-mannosidase (Bt-α1,3-MNS) (**Figure 2C and D**), respectively, suggesting that the rituximab N-glycans are less accessible. Therefore, we used higher enzyme amounts and longer incubation times to acquire a full conversion in the consecutive enzymatic steps on rituximab (**Table 2**).

In the natural N-glycan processing pathway, Man₅GlcNAc₂ glycans are modified in the Golgi apparatus by the GNT-I enzyme, which adds a GlcNAc to the α1,3-linked mannose arm (**Figure 1A**). We demonstrated that the RnGNT-I enzyme can similarly modify Man₅GlcNAc₂ to form GlcNAcMan₅GlcNAc₂ (**Figure S6**). In the Golgi, GlcNAcMan₅GlcNAc₂ is further trimmed down to GlcNAcMan₃GlcNAc₂ by Golgi α-mannosidase II, which can also be performed *in vitro* with the bacterial Bt-α1,6-MNS and Bt-α1,3-MNS enzymes (**Figure S6**). Though our enzymes can replicate this natural glycan processing pathway, it is not the most efficient strategy to generate GlcNAcMan_3_GlcNAc_2_ *in vitro*. The mannosidases Ck-α1,2-MNS, Bt-α1,6-MNS, and Bt-α1,3-MNS can also be applied together to generate Man₃GlcNAc₂ in a one-pot reaction (**Figure S7**). The resulting Man_3_GlcNAc_2_ substrate can be modified by RnGNT-I to generate GlcNAcMan_2_GlcNAc_2_ as well (**Figure 2E**). Thus, in contrast to the natural processing sequence, the most efficient way to generate GlcNAcMan₃GlcNAc₂ glycans *in vitro* is to first trim down the mannoses, and then build up the GlcNAc with RnGNT-I. The GlcNAcMan₃GlcNAc₂ substrate can then be further modified by HsGNT-II to generate a GlcNAc2Man₃GlcNAc₂ (**Figure 2F**), which enables the *in vitro* enzymatic pathway to complex glycans.

Next, we tried to modify single and biantennary N-glycans with terminal GlcNAc to a core-fucosylated form with our MmFUT8 enzyme. Although the *in vitro* modification occurred, it was only observed in low levels (**Figure S8**). OoASP-1 received approximately 40% core α1,6-fucose on the single antenna form, while the double antenna glycoform only received 5-10% fucose. Rituximab hardly received any core α1,6-fucose, emphasizing the more hidden nature of the N-glycans. The MmFUT8 was only purifiable in low levels and was only visible on an anti-HIS western blot (**Figure S4**), which could be the cause for the low levels of core-fucosylation in these *in vitro* reactions.

Non-fucosylated glycoforms were further modified with the MmGALT1 enzyme (**Figure 3**). We were able to fully galactosylate the GlcNAcMan₃GlcNAc₂ glycan on both OoASP-1 and rituximab. In parallel, we successfully modified GlcNAcMan₅GlcNAc₂ and GlcNAcMan₄GlcNAc₂ with a terminal galactose on Oo-ASP1. These glycans do not naturally occur due to the sequential processing steps of the Golgi apparatus, but could be generated *in vitro* (**Figure S9**). The biantennary GlcNAc_2_Man_3_GlcNAc₂ N-glycans were fully galactosylated on OoASP-1 by the MBP-tagged MmGALT1 (**Figure 3B**, left), whereas the enzyme was less efficient on the N-glycans of rituximab (**Figure 3B**, right). Glycans in the Fc portion of the IgG antibody have a more inward confirmation, which could possibly affect the accessibility of the N-glycan by glycoenzymes [58]. We hypothesized that the relatively large size of the MBP-tag (42 kDa) could hamper with the glycan accessibility, which is why we performed *in vitro* modifications with a Small Ubiquitin-like Modifier (SUMO, 11 kDa) tagged version of the MmGALT1 instead. This version was able to fully galactosylate the biantennary N-glycans on rituximab (**Figure 3C**), strengthening our hypothesis.

**Figure 3.**
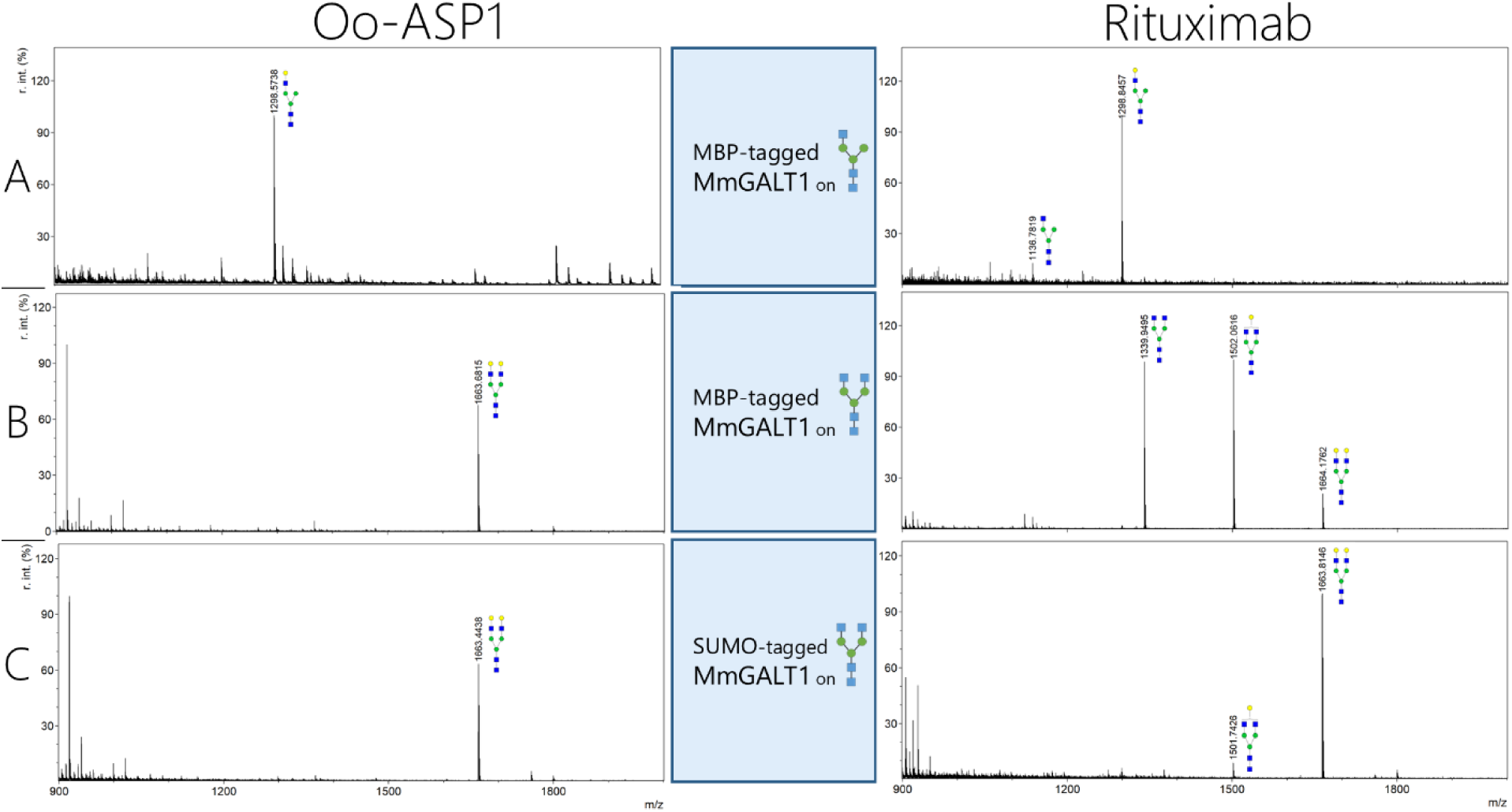
MALDI-TOF MS spectra of PNGase F released glycans generated on OoASP-1 and Rituximab through *in vitro* galactosylation by SUMO and MBP-tagged MmGALT1. *In vitro* glycoengineered OoASP-1 and Rituximab, containing GlcNAcMan₃GlcNAc₂ and GlcNAcMan₅GlcNAc₂ glycans, were galactosylated by SUMO and MBP-tagged MmGALT1 as depicted in the blue boxes. Peaks that could not be labeled to glycans are left unannotated. Glycan structures are drawn according to symbol nomenclature for glycans (SNFG).

Thus far, our *in vitro* system was not successful in generating all envisioned glycoforms, which is mainly due to the low activity of MmFUT8 on our glycoprotein substrates. *In vitro* glycosylation steps are typically time-consuming and costly, primarily due to the high costs of UDP- and GDP-linked sugar donors. To address this, we proceeded to apply a plant-based glycoengineering approach to directly generate some of the N-glycans that would otherwise require multiple *in vitro* engineering steps or are difficult to obtain *in vitro*. These substrates could then serve as a starting point for *in vitro* glycoengineering to achieve the pharmaceutically relevant target glycoforms.

### *In planta* glycoengineering of OoASP-1 and rituximab

To produce OoASP-1 and rituximab with various N-glycan structures *in planta*, kifunensine was omitted from the infiltration suspension and instead several glycosyltransferases or glycosidases were co-expressed. Without infiltrating any glycoenzyme constructs or kifunensine, OoASP-1 and rituximab receive relatively homogeneous GlcNAc_2_Man_3_GlcNAc_2_ glycans upon expression in *N. benthamiana* (**Figure 4A**), creating a good starting point for the build-up of complex glycans *in vitro*. To generate core α1,6-fucosylated complex N-glycans, DmFut8 was co-infiltrated. This yields fully homogeneous GlcNAc_2_Man_3_GlcNAc_2_Fuc glycans on rituximab, but yields some heterogeneity regarding fucosylation and more truncated N-glycans on OoASP-1 (**Figure 4B**). Glycoengineering of *N. benthamiana* could also be applied to directly generate Man_3_GlcNAc_2_ and Man_5_GlcNAc_2_ glycans through co-expression of NbHEXO3 or a combination NbHEXO3, DrGALT and NbBGAL8, respectively (strategy previously proposed by van der Kaaij et al. [28]). Here, engineering of Man_3_GlcNAc_2_ was much more homogeneous on OoASP-1 than on rituximab (**Figure 4C**), whereas Man_5_GlcNAc_2_ engineering showed similar levels of homogeneity on both substrates (**Figure 4D**). Direct engineering of hybrid GlcNAcMan_5_GlcNAc_2_ glycans was also attempted through co-infiltration of DrGALT and NbBGAL8. This resulted in relatively homogeneous N-glycans on rituximab, but yielded a heterogeneous N-glycan composition on OoASP-1 (**Figure 4E**). These results show that plant glycoengineering can be used to generate a variety of glycoforms with less effort compared to *in vitro* engineering, but that N-glycan homogeneity is affected by protein substrate.

**Figure 4.**
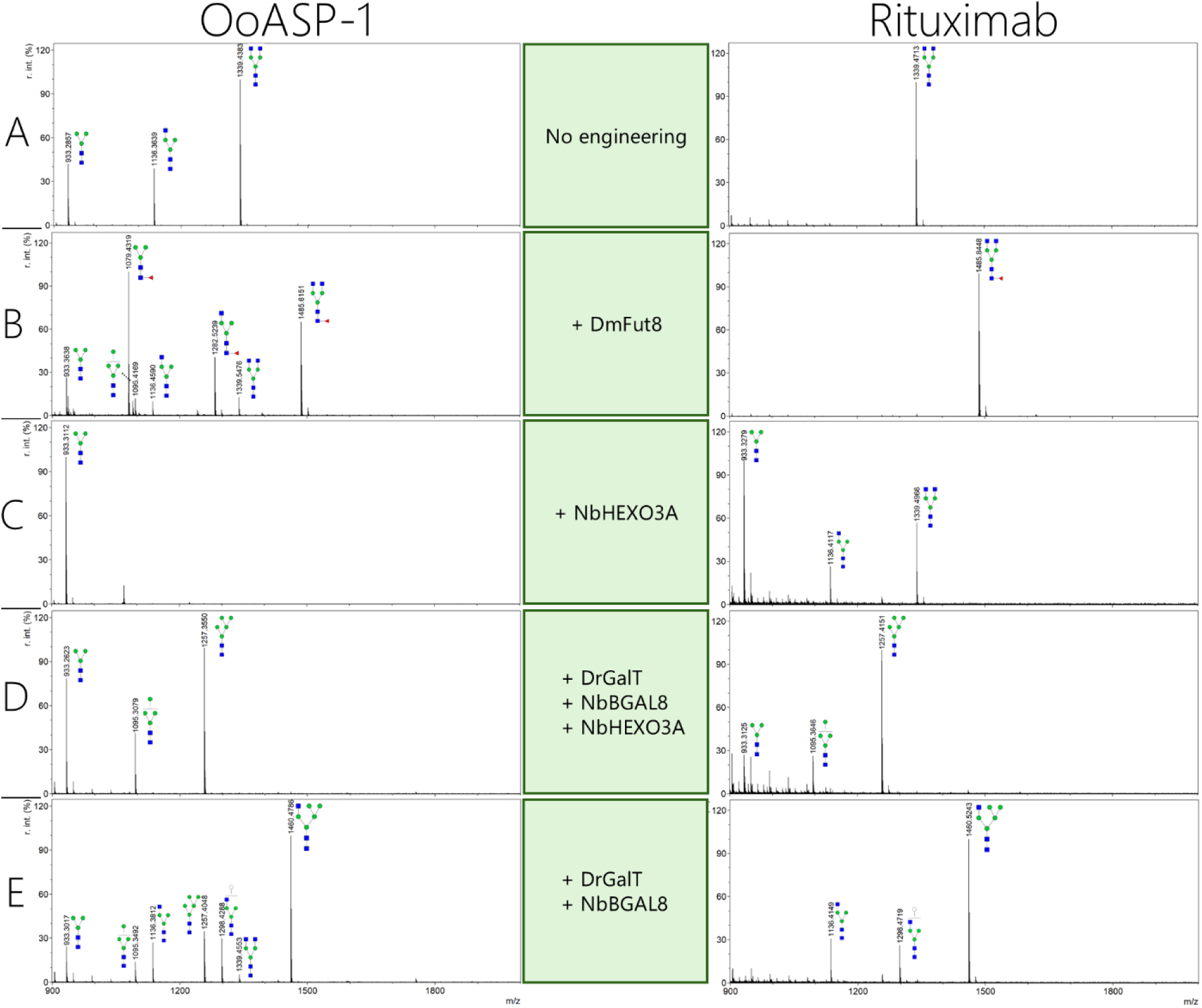
MALDI-TOF MS spectra of PNGase F released glycans generated on OoASP-1 and rituximab through *in planta* glycoengineering. *N. benthamiana* plants were infiltrated with *A. tumefaciens* strains harboring expression constructs for OoASP-1 (left column) or rituximab (right column), together with strains harboring expression constructs for relevant glycosyltransferases or glycosidases (depicted in middle column). Each row (**A-E**) represents a different glycoengineering strategy. Peaks that could not be labeled to glycans are left unannotated. Glycan structures are drawn according to symbol nomenclature for glycans (SNFG).

### Combining *in planta* and *in vitro* glycoengineering strategies to generate pharmaceutically relevant N-glycans on OoASP-1 and rituximab

Next, we sought to efficiently generate all pharmaceutically relevant glycoforms of OoASP-1 and rituximab by combining *in planta* and *in vitro* engineering strategies. OoASP-1 naturally carries predominantly GalGlcNAcMan_3_GlcNAc_2_Fuc, and to a lesser extent Gal_2_GlcNAc_2_Man_3_GlcNAc_2_Fuc, GlcNAcMan_3_GlcNAc_2_Fuc and Man_3_GlcNAc_2_Fuc glycans [6]. On all these N-glycans, about three-quarters of core fucoses are α1,6-linked, and one quarter is α1,3-linked [6].

To create the α1,6-fucosylated glycoforms, OoASP-1 was first produced in *N. benthamiana* with co-expression of NbHEXO3A and DmFUT8. In contrast to the heterogeneous N-glycans found on OoASP-1 upon co-expression of DmFUT8 only (**Figure 4B**), the combination of NbHEXO3A and DmFUT8 yielded Man_3_GlcNAc_2_Fuc glycans with minimal heterogeneity (**Figure 5A**). The N-glycan on OoASP-1 was further modified *in vitro* with RnGNT-I, HsGNT-II and MmB4GALT to generate GlcNAcMan_3_GlcNAc_2_Fuc (**Figure 5B**), GalGlcNAcMan_3_GlcNAc_2_Fuc (**Figure 5C**) and Gal_2_GlcNAc_2_Man_3_GlcNAc_2_Fuc (**Figure 5D**).

**Figure 5.**
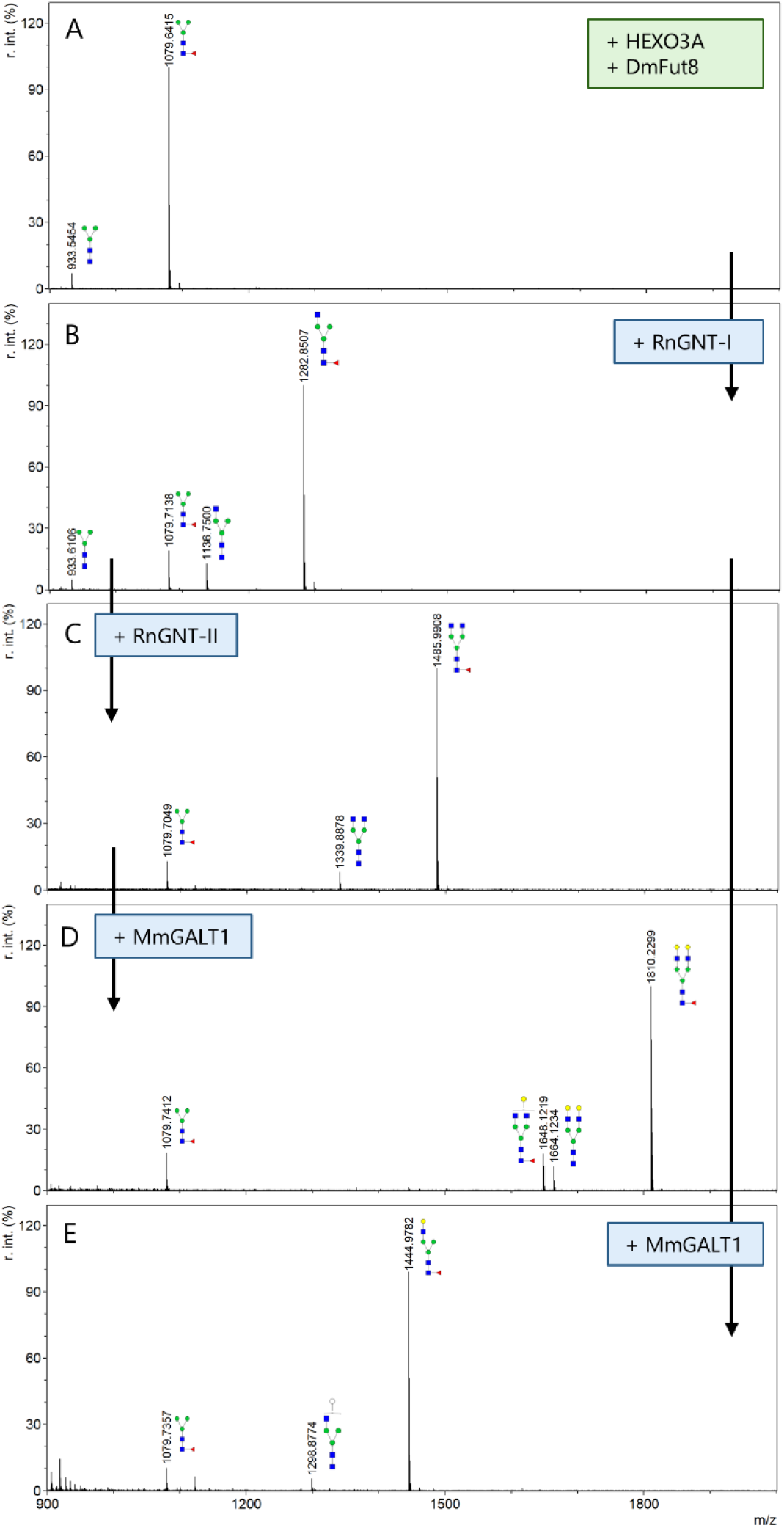
MALDI-TOF MS spectra of PNGase F glycans generated on OoASP-1 through combining *in planta* and *in vitro* glycoengineering. *N. benthamiana* plants were infiltrated with *A. tumefaciens* strains harboring expression constructs for OoASP-1, NbHEXO3A and DmFUT8 to obtain OoASP-1 with Man_3_GlcNAc_2_Fuc glycans (**A**). Subsequent *in vitro* glycoengineering with RnGNT-I, RnGNT-II and MmGALT1 was performed to generate pharmaceutically relevant native OoASP-1 glycans (**B, C, D, E**). Green boxes depict constructs used for *in planta* glycoengineering, blue boxes depict enzymes used for *in vitro* glycoengineering. Black arrows indicate sequential *in vitro* reactions. Peaks that could not be labeled to glycans are left unannotated. Glycan structures are drawn according to symbol nomenclature for glycans (SNFG).

To generate OoASP-1 glycoforms with α1,3-core fucosylated glycans, the protein was produced *in planta* with Man_3_GlcNAc_2_Fuc glycans through co-expression of NbHEXO3A and SmFucTC (**Figure S10**). However, further *in vitro* modification yielded N-glycans with low homogeneity, possibly due to the limited capacity of GNT-I to modify the α1,3-fucosylated substrate (**Figure S10**). The N-glycans described in figure 5C, D and E could also be generated on OoASP-1 without core fucose with high homogeneity through omitting DmFUT8 from the infiltration suspension (**Figure S11**).

Rituximab is currently produced and marketed as a heterogeneously glycosylated antibody, predominantly bearing core α1,6-fucosylated complex N-glycans with low to moderate terminal galactosylation [59]. Structural and functional analyses show that galactosylation at the α6-antenna of the Fc N-glycan can increase FcγRIIIa (CD16A) binding affinity and thereby enhance antibody-dependent cellular cytotoxicity (ADCC). This effect is reduced by core α1,6-fucose, which sterically hinders FcγRIIIa engagement [60]. While core fucosylation is less beneficial for a potent ADCC response, it can be beneficial in the context where excessive immune activation is undesirable and may help to mitigate risks associated with overactive immune responses [61,62]. Therefore, we focused on making both fucosylated and afucosylated glycoforms of rituximab with terminal galactose, since both variants could be important for tailored treatment of patients. We were able to directly produce homogeneous GlcNAc_2_Man_3_GlcNAc_2_ glycans *in planta* with or without core α1,6-linked fucose (**Figure 6A and C**). These antibody glycoforms were then further modified *in vitro* with SUMO-tagged MmGALT1 to generate homogeneously galactosylated diantennary N-glycans (**Figure 6B and D**). Taken together, these results show that by combining *in planta* and *in vitro* engineering strategies, we were able to homogeneously generate a wide variety of glycoforms (**Figure 1B**) on both OoASP-1 and rituximab with limited engineering efforts.

**Figure 6.**
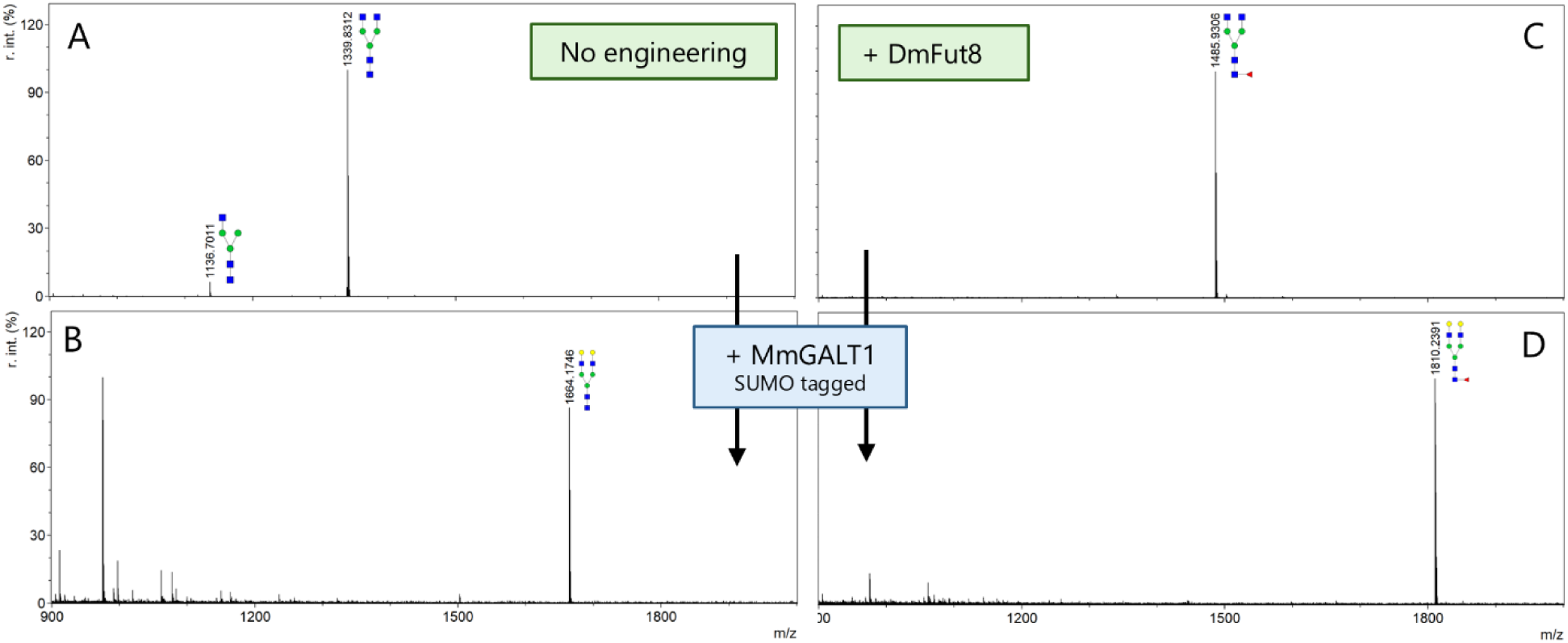
MALDI-TOF MS spectra of glycans generated on rituximab through combining *in planta* and *in vitro* glycoengineering. *N. benthamiana* plants were infiltrated with *A. tumefaciens* strains harboring expression constructs for rituximab and DmFut8 to obtain GlcNAc_2_Man_3_GlcNAc_2_ and GlcNAc_2_Man_3_GlcNAc_2_Fuc glycans (**A** and **B**, respectively). Both substrates were then modified *in vitro* with SUMO-tagged HsGALT1 to facilitate galactosylation (**C** and **D**). Green boxes depict constructs used for *in planta* glycoengineering, blue boxes depict enzymes used for *in vitro* glycoengineering. Black arrows indicate sequential *in vitro* reactions. Peaks that could not be labeled to glycans are left unannotated. Glycan structures are drawn according to symbol nomenclature for glycans (SNFG).

## Discussion

The objective of this study was to develop a versatile production platform for the quick and resource efficient generation of homogeneously glycosylated biopharmaceuticals. Our results demonstrate that the combination of *in planta* and *in vitro* glycoengineering enables the production of target proteins decorated with a wide range of N-glycans, including high mannose, paucimannose, hybrid, and complex types. In addition, it could generate glycoforms that are of pharmaceutical relevance on a monoclonal antibody and a vaccine candidate, but is most likely applicable on a wide range of target proteins.

Compared to previously published *in vitro* glycoengineering systems, our enzyme purification methods are on par, without relying solely on large solubility enhancing tags [36,45]. Jaroentomeechai et al. [36] made a universal pipeline to synthesize a broad range of glycoenzymes by utilizing the solubilizing enhancing N-terminal MBP-tag and a C-terminal truncated version of the human apolipoprotein A1, which drastically enhanced the solubility of the enzymes they tried to produce in *E. coli*. Although it worked to produce and purify a wide range of active glycoenzymes, there is a risk that dual tags result in an inactive protein or affect the accessibility of the enzyme, as we have demonstrated with our MBP-tagged version of the MmGALT1 on rituximab. Similarly, Zhang et al. [45] produced a wide range of glycoenzymes, most of which received an N-terminal MBP-tag, but still resulted in a low solubility of their HsGNT-I and HsGNT-II enzymes. Our work shows that smaller tags, such as SUMO (11 kDa) and thioredoxin (12 kDa), can achieve solubility levels comparable to the much larger MBP tag (42 kDa). In the future, it would be advisable to produce insoluble or less soluble glycoenzymes with a variety of solubility-enhancing tags, as it is difficult to predict how a specific tag will affect the enzyme’s solubility and activity.

In this study we demonstrate that our *in vitro* glycoengineering system efficiently modifies a broad spectrum of glycoforms, from high mannose to complex structures, using a universal buffer and minimal amounts of costly donor sugars. Our reactions were able to yield homogeneous glycoforms with enzyme usage comparable to previous studies, yet require significantly less donor sugar in most reactions (**Table 2 & Table S4**). Since donor sugar prices can go up to a few thousand US dollars per gram, it is more cost-efficient to perform *in vitro* engineering reactions at lower donor sugar concentrations.

Although we could synthesize the majority of the envisioned N-glycans *in vitro*, we were not able to synthesize homogeneously core-fucosylated N-glycans on OoASP-1 and rituximab *in vitro*. FUT8 is difficult to produce in *E. coli*, as shown here and by Jaroentomeechai et al. [36], but can be readily expressed in large amounts in human HEK293 cells [63]. MmFUT8 contains four predicted disulfide bonds but may also require other post-translational modifications, such as phosphorylation at Ser278 [64]. In addition, many eukaryotic proteins require chaperones for proper folding, but identifying the correct functional eukaryotic chaperone for MmFUT8 production in *E. coli* can be challenging and time-consuming [65]. The limited *in vitro* incorporation of α1,6-fucose could be addressed by glycoengineering the enzymatic machinery for core α1,6-fucosylation into *N. benthamiana*, thereby reducing the need for the lengthy and costly process of optimizing *in vitro* glycoengineering. As such, the combination of *in planta* and *in vitro* glycoengineering utilizes the benefits of both engineering systems to their fullest. Smart combinations of strategies can reduce the amount of *in vitro* engineering steps from six (for Gal_2_GlcNAc_2_Man_3_GlcNAc_2_ on rituximab) to just one, while incorporating fucose on a level that would not have been possible *in vitro*.

We also show the application of several *in planta* glycoengineering strategies that were previously proposed, but not all executed, by van der Kaaij et al. [28]. Our results indicate that the proposed strategies are feasible, but with varying levels of homogeneity. For example, the engineering strategies applied for the generation of GlcNAcMan_5_GlcNAc_2_ and Man_5_GlcNAc_2_ yield relatively heterogeneous N-glycans. The two engineering strategies were devised based on previous studies showing the generation of GalGlcNAcMan_5_GlcNAc_2_ through overexpression of 35S:sialDrGalT. In theory, addition of a β-galactosidase and β-hexosaminidase should result in trimming of this N-glycan to GlcNAcMan_5_GlcNAc_2_ and Man_5_GlcNAc_2_ respectively, but our results show that this does not yield homogeneous N-glycans. The level of heterogeneity can be explained by the complex nature of *in vivo* N-glycosylation. N-glycan processing in the Golgi is a sequential process, and therefore the sub-localization of glycoenzymes impacts the result of glycoengineering [66]. Though the exact mechanism determining Golgi sub-localization of glycan processing enzymes remains unknown, is it widely accepted that the enzymes’ transmembrane domain (TMD) plays a major role in determining its localization [66]. Thus, glycan processing can be optimized by targeting transferases to specific Golgi cisternae through the generation of chimeric glycosyltransferases (carrying non-native TMDs) [47,67]. Additionally, promoter strength can also influence the efficiency of glycoengineering [47,68]. Finetuning promoter strength and Golgi localization could possibly optimize *in planta* generation of GlcNAcMan_5_GlcNAc_2_ and Man_5_GlcNAc_2_ but, at this point, *in vitro* glycoengineering remains the best strategy to homogeneously generate these N-glycans.

In contrast, the generation of Man_3_GlcNAc_2_, GlcNAc_2_Man_3_GlcNAc_2_ and fucosylated N-glycans can be greatly simplified through the application of *in planta* glycoengineering. Both OoASP-1 and rituximab could be produced *in planta* with near complete fucosylated Man_3_GlcNAc_2_ and GlcNAc_2_Man_3_GlcNAc_2_ glycans, respectively. Though both substrates can be fucosylated efficiently, the efficiency of Man_3_GlcNAc_2_ and GlcNAc_2_Man_3_GlcNAc_2_ generation varies between OoASP-1 and rituximab. This variability can be explained by differences in N-glycan location within the protein backbone. The location of a N-glycan determines its accessibility to glycoenzymes, and consequently impacts the level to which a N-glycan is processed [69,70]. The N-glycan on OoASP-1 is exposed on the surface of the protein and can therefore readily be accessed by glycoenzymes [6]. In contrast, the N-glycan of rituximab is located in the partially shielded Fc region, a location which has previously been reported to be aberrantly glycosylated in plants [71,72]. We show that without any glycoengineering in *N. benthamiana*, the terminal GlcNAc residues on OoASP-1 are partially trimmed off, likely due to the presence of hexosaminidases in the apoplast [48]. In contrast, the rituximab N-glycan remains fully homogeneous, as it is likely shielded from hexosaminidase activity. The same mechanism explains why the *in vivo* engineering of Man_3_GlcNAc_2_ on OoASP-1 is fully efficient upon co-expression of HEXO3A, whereas rituximab retains a fraction of GlcNAc_2_Man_3_GlcNAc_2_ glycans. Thus, though *in planta* glycoengineering can produce fully homogeneous N-glycans, the best engineering strategy will always depend on the intrinsic characteristics of each target protein.

In the *in vitro* glycoengineering of rituximab and OoASP-1, we observed similar effects of protein characteristics on glycoengineering efficacy. Rituximab required significantly longer incubation steps and more donor sugar to be fully converted to the desired glycoforms compared to OoASP-1. In addition, both OoASP-1 and SmKappa-5 (previously reported by van der Kaaij et al [31]) could be fully galactosylated with the MBP-tagged MmGALT1, while rituximab required the smaller SUMO-tagged variant to be fully galactosylated. Terminal galactosylation on the α6 antenna of Fc N-glycans in IgG antibodies alters the glycan conformation, partially burying it within the Fc region [60]. This likely explains why, in most commercially available rituximab, galactosylation occurs predominantly on a single antenna [59] and why *in vitro* glycoengineering could help to make a more potent rituximab.

In the future, the precision glycoengineering system presented here could be further advanced by expanding its N-glycan repertoire. We have not yet included strategies for the generation of sialylated, multi-antennary or bisecting N-glycans, though they are of therapeutic interest. Of the three, sialic acid is the most well studied in the context of pharmaceuticals. Presence of sialic acid can influence *in vivo* half-life of pharmaceuticals [73,74] or influence immune responses due to differential interaction with Fc [75] or lectin receptors on leukocytes [76]. Sialic acids on vertebrate N-glycans are typically attached to galactose via an α2,3 or α2,6 linkage, but can also be attached to another sialic acid via an α2,8 linkage to form polysialic acid (PSA) [77]. For all linkage types, both *in planta* and *in vitro* glycoengineering strategies are described. Multiple studies have succeeded to produce α2,6-sialylated N-glycans in *N. benthamiana* [78–81]. Kallolimath et al [78] also generated glycoproteins in plants that carry only α2,3 linked sialic acid. Since plants do not naturally decorate their N-glycans with sialic acid, production of sialylated N-glycans requires introduction of the complete biosynthetic pathway compromising 6 enzymes [82]. This makes it possible to tune the introduced linkage type by swapping the type of sialyltransferase that is introduced [78]. Nonetheless, sialylation in *Nicotiana benthamiana* remains heterogeneous and thus requires further optimization to enable the production of uniformly sialylated proteins for structure–function analyses. Therefore, the engineering system presented here might best be elaborated with *in vitro* incorporation of sialic acid. Several studies already have homogeneously incorporated sialic acid on N-glycans in α2,3-, α2,6- and α2,8-linkages through *in vitro* enzymatic sialylation [33,36,83,84] demonstrating the feasibility of adding sialic acid to the N-glycan repertoire presented here.

In contrast, homogeneous incorporation of multi-antennary and bisecting N-glycans might be more complicated. While they are known to play a role in several diseases - such Alzheimer’s disease [85,86] and breast cancer [87,88] - their known effects on biopharmaceuticals remain limited to increasing the half-life of EPO [74,89]. Multiple studies report the introduction of chimeric GNT-III or GNTI-IV and GNT-V in *N. benthamiana* for generating bisecting or tri- and tetra-antennary glycans, respectively [90–93]. Though the expression of single glycosyltransferases leads to impressive increases in N-glycan branching, it is not yet homogeneous to the level desired for structure-function studies. In this case, *in vitro* glycoengineering offers no straightforward alternative, as, to our knowledge, no studies have yet applied *in vitro* glycoengineering to produce multi-antennary or bisecting N-glycans on glycoproteins. However, Hamilton et al. [94] show enzymatic synthesis of bisecting or multi-antennary free N-glycans using commercially available *Homo sapiens* GNT-III and -V, and *E. coli* produced *Bos taurus* GNT-IV. Though the Golgi-localized forms of these enzymes naturally modify glycoproteins, it remains to be determined whether their recombinant soluble forms retain their function on protein-linked glycans in an *in vitro* setting.

Beyond its application for structure-function relationships, the glycoengineering system presented here could, upon further optimization, evolve into a large-scale production system for homogeneous biopharmaceuticals. Biopharmaceuticals or glycoenzymes can be immobilized to enable efficient sequential *in vitro* glycan modification without cross-mixing of the reaction components after the final purification steps. For example, Jaroentomeechai et al.[36] immobilized IgG antibodies on protein A/G beads, allowing stepwise enzymatic modification of the antibody, without enzymatic leaching in the end product. Another approach is to immobilize the glycoenzymes, enabling reaction compartmentalization, enhanced enzymatic activity, improve reusability, and prevent substrate– enzyme interaction after the final steps [95,96]. This could be turned into a flow-based system as demonstrated by Aquino et al, 2021 [95], but will require a lengthy optimization process. Moreover, this approach could become cost-effective if sugar donors can be produced more cheaply in the future.

The combination of such a production grade *in vitro* glycoengineering pipeline with protein production in *N. benthamiana* strengthens its potential as a biopharmaceutical production system. Protein production in *N. benthamiana* is fast, cheap, scalable, and is increasingly being put forward as an alternative to mammalian expression systems in the context of pandemic-preparedness [97–99]. Though market approval of plant-produced pharmaceuticals can be faced with regulatory hurdles [100], precedent of Protalix’ carrot cell produced taliglucerase alfa [101] and Medicago’s *N. benthamiana* produced COVID-19 vaccine [102] shows that it is feasible. As such, further development of the glycoengineering system presented here could not only benefit our knowledge of structure-function relationships of N-glycans, but also contribute to high quality biopharmaceutical production in pandemic-response situations.

## Supporting information

Supplemental Table S1: Nucleotide Sequences of Rituximab Heavy and Light Chains produced in this study.

Supplemental Table S2: Codon-Optimized Synthetic Enzyme Genes Expressed in *E. coli*.

Supplemental Table S3 : Primers used in this study.

Supplementary Table S4: Summary of Enzymes, Conditions, and Substrates Used in Previous Glycosylation Studies

Supplemental figure S1: Coomassie Blue stained SDS-PAGE gels of purification of high mannose Oo-ASP1 and Rituximab

Supplemental Figure S2: Coomassie Brilliant Blue-stained SDS-PAGE gels showing purification of RnGNT-I and MmGALT1, with and without solubility-enhancing tags.

Supplemental Figure S3: Coomassie Brilliant Blue-stained SDS-PAGE gels showing purification of RnGNT-I, MmGALT1, Ck-α1,2-MNS, Bt-α1,3-MNS, Bt-α1,6-MNS.

Supplemental Figure S4 : Coomassie Brilliant Blue-stained SDS-PAGE gels and a western blot showing the purification of MmFUT8 and HsGNT-II

Supplementary Figure S5: MALDI-TOF MS spectra of glycans generated on rituximab through sequential in vitro incubation with mannosidases in varying ratios.

Supplementary Figure S6: MALDI-TOF MS spectra of GlcNAcMan_3_GlcNAc_2_ glycans generated on OoASP-1 and rituximab via the natural glycan processing route.

Supplementary figure S7: MALDI-TOF MS spectra of Man_3_GlcNAc_2_ glycans generated on OoASP-1 and rituximab in a one-pot reaction.

Supplementary Figure S8: MALDI-TOF MS spectra of *in vitro* fucosylated glycans generated on rituximab.

Supplementary Figure S9: MALDI-TOF MS spectra of unusual galactosylated glycans generated on OoASP-1.

Supplementary Figure S10. MALDI-TOF MS spectra of PNGase L released glycans generated on OoASP-1 through combining *in planta* and *in vitro* glycoengineering.

Supplementary Figure S11. MALDI-TOF MS spectra of PNGase F released glycans generated on OoASP-1 through combining *in planta* and *in vitro* glycoengineering.

## Declaration of generative AI and AI-assisted technologies in the writing process

During the preparation of this work the author(s) used Consensus and ChatGPT as a search engine and to provide suggestions of optimization of individual paragraph or sentence structure. After using this tool/service, the author(s) reviewed and edited the content as needed and take(s) full responsibility for the content of the published article.

## Data availability

All data underlying the manuscript will be made available upon reasonable request.

## Authorship statement

PN, AS and RHPW conceived the project. RHPW and AS acquired funding. MRS and PN designed and performed the experiments, advised by AS and RHPW. MRS and PN analyzed the data. MRS and PN wrote and revised the draft manuscript that was edited by AS and RHPW. All authors read and approved the final article.

## Acknowledgements

This research was performed within Eurostars project 115017 ‘HiT-GlyP’ and PPS ‘Wise with Worms’ (LWV22248, BO-63-001-066). Wise with Worms co-financed by the Top Consortium for Knowledge and Innovation ‘Agri & Food’ by the Dutch Ministry of Agriculture, Fisheries, Food Security and Nature. The project was supported by Royal GD, LTO Nederland, zLTO, LTO Noord, COV, the provinces: North-Holland, South-Holland, Utrecht, Overijssel, Friesland and Groningen and the commercial partner Zoetis. We would like to thank Bo Briggeman, Kostas Gaitanis, Leon ten Hoopen and Magdalini Koroxenidou for their contributions to the practical work of this study.

## Notes

### Competing Interest Statement

The authors have declared no competing interest.

