## Supplemental Figures for "Precision Glycoform Engineering: Combining plant and *in vitro* systems for tailored biopharmaceutical production"

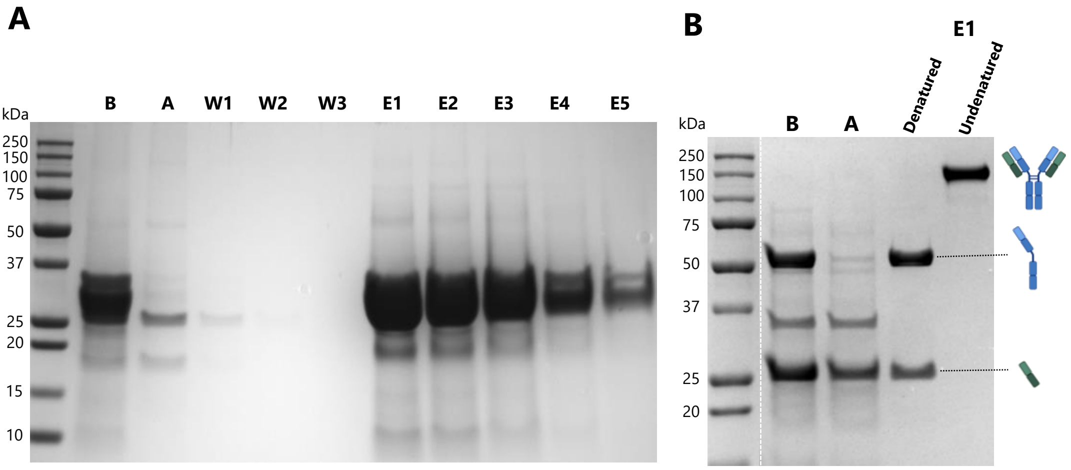


**Supplemental Figure S1: Coomassie Blue stained SDS-PAGE gels of purification of high mannose Oo-ASP1 and Rituximab**

**(A)** Cation exchange resin purification of high mannose Oo-ASP1

**(B)** The purification of High mannose Rituximab with protein A

**B** = Before binding to resin, **A** = After binding to resin, **W**= Wash, **E** = Elution.

Oo-ASP1 (glycosylated) = ≈27-32 kDa,, Rituximab = 150 kDa, **γ**= 50 kDa, **κ**= 25 kDa

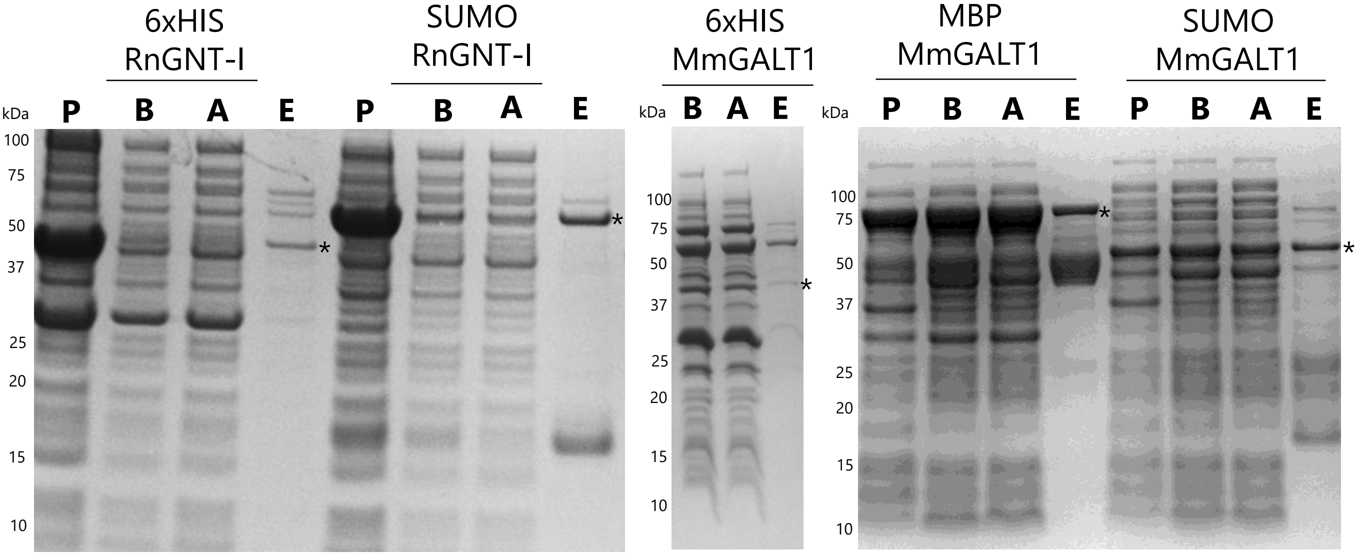


**Supplemental Figure S2: Coomassie Brilliant Blue-stained SDS-PAGE gels showing purification of RnGNT-I and MmGALT1, with and without solubility-enhancing tags.**

The glycoenzymes were expressed in BL21 *E. coli* cells and purified with HisPur™ Ni-NTA Resin. The subsequent purification steps are visualized with SDS-PAGE and Coomassie blue staining. **P** = Pellet solubilized in 5% SDS buffer, **B** = Before binding to the Ni-NTA resin, **A** = After binding to the Ni-NTA resin, **E** = Elution, * = protein of interest

6xHIS-RnGNT-I = 48.86 kDa, 6xHIS-SUMO-RnGNT-I = 60.34 kDa, 6xHIS-MmGALT1 = 41.18 kDa, SUMO-6xHIS-MmGALT1 = 52.66 kDa, MBP-6xHIS-MmGALT1 = 81.63 kDa

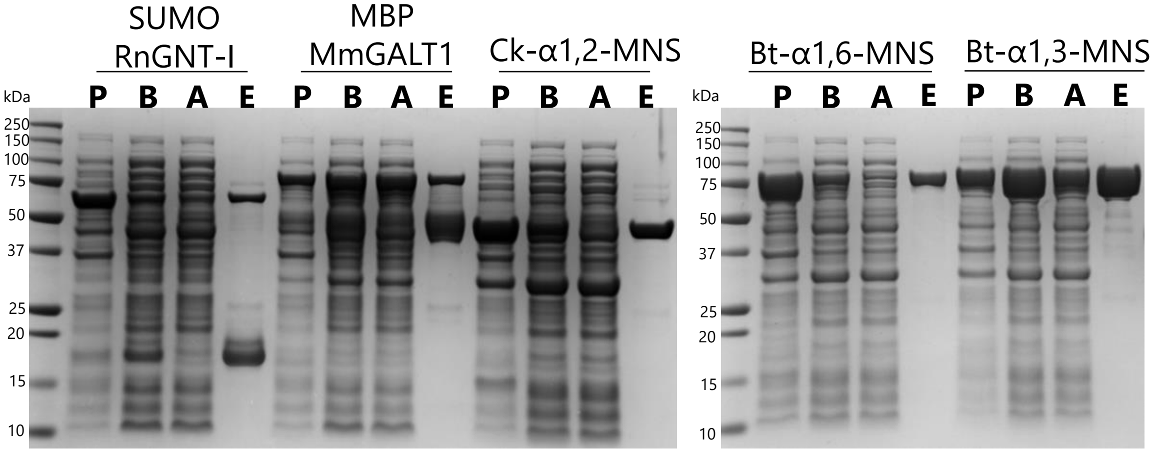


**Supplemental Figure S3: Coomassie Brilliant Blue-stained SDS-PAGE gels showing purification of RnGNT-I, MmGALT1, Ck-α1,2-MNS, Bt-α1,3-MNS, Bt-α1,6-MNS**

The glycoenzymes were expressed in BL21 *E. coli* cells and purified with HisPur™ Ni-NTA Resin. The subsequent purification steps are visualized with SDS-PAGE and Coomassie blue staining.

**P** = Pellet solubilized in 5% SDS buffer, **B** = Before binding to the Ni-NTA resin, **A** = After binding to the Ni-NTA resin, **E** = Elution.

SUMO-6xHIS-RnGNT-I = 60.34 kDa, MBP-6xHIS-MmGALT1 = 81.63 kDa, 6xHIS-CkGH47 = 50.57 kDa, 6xHIS-Bt3994 = 83.1 kDa, 6xHIS-Bt1769 = 84.02 kDa.

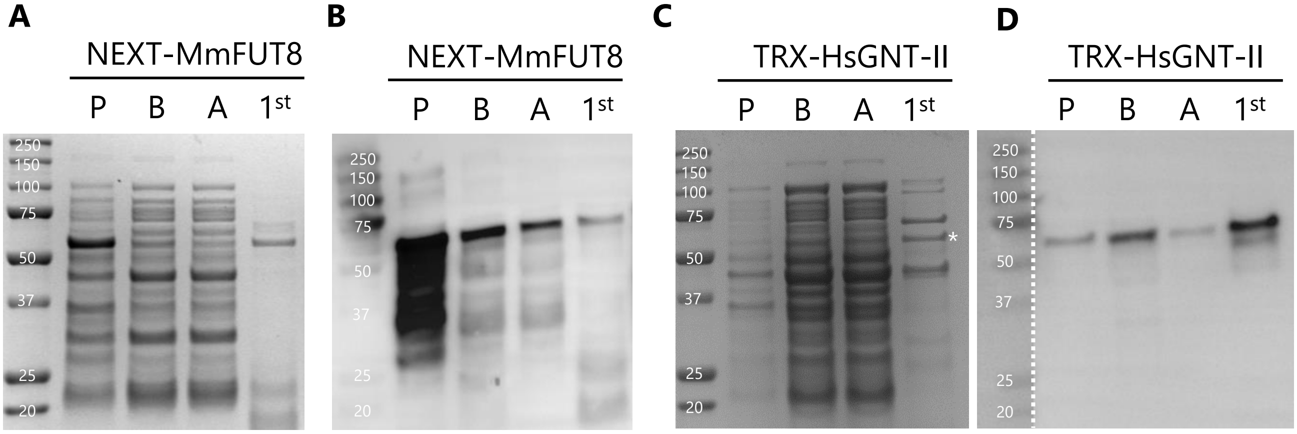


**Supplemental Figure S4 : Coomassie Brilliant Blue-stained SDS-PAGE gels and a western blot showing the purification of MmFUT8 and HsGNT-II**

MmFUT8 and HsGNT-II were expressed in Rosetta-Gami 2 (DE3)pLysS cells and purified with HisPur™ Ni-NTA Resin. The subsequent purification steps are visualized with SDS-PAGE and Coomassie blue staining **(A + C)** and an anti-HIS tag western blot **(B + D).**

**P** = Pellet solubilized in 5% SDS buffer, **B** = Before binding to the Ni-NTA resin, **A** = After binding to the Ni-NTA resin, **E** = Elution. MmFUT8 = 61.17 kDa , HsGNT-II = 60.4 kDa, * indicates TRX-HsGNT-II

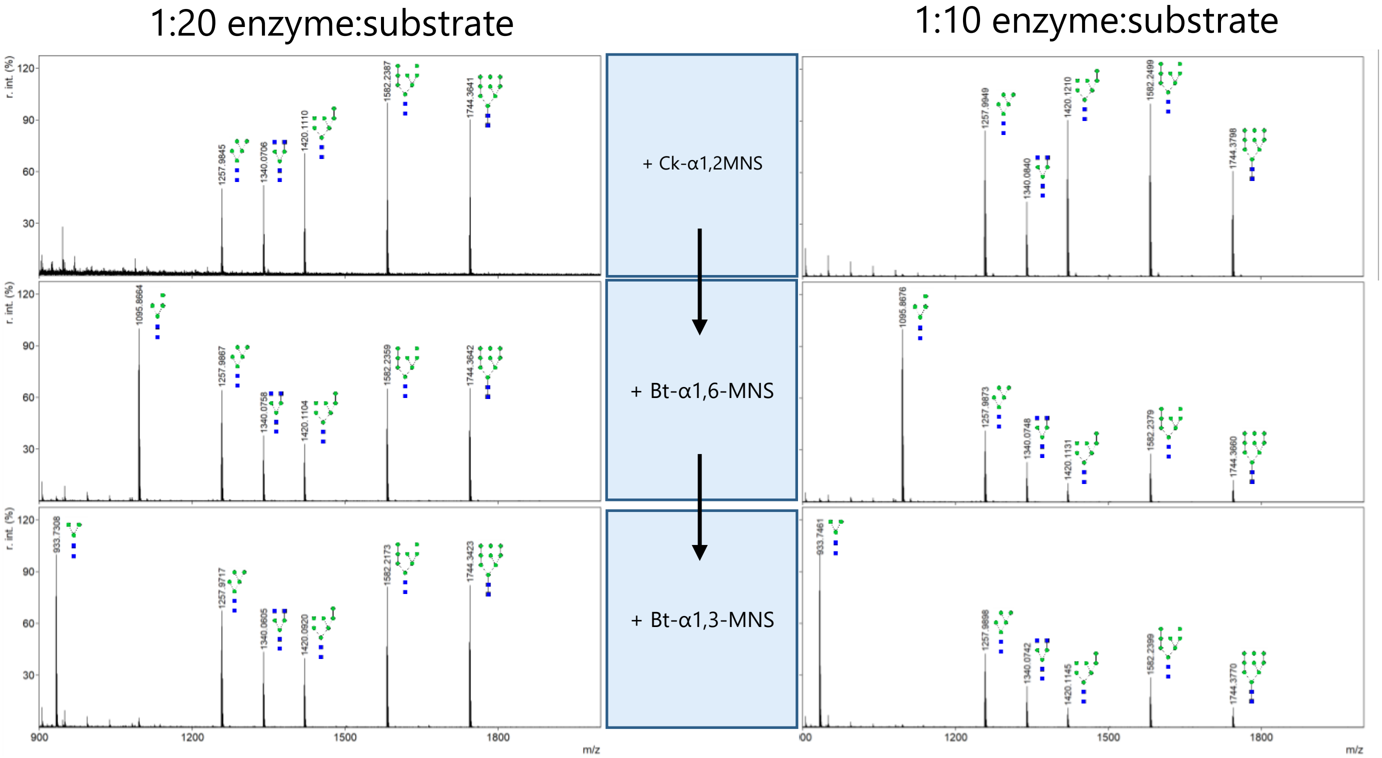


**Supplementary Figure S5: MALDI-TOF MS spectra of glycans generated on rituximab through sequential in vitro incubation with mannosidases in varying ratios.** *N. benthamiana* plants were infiltrated with *A. tumefaciens* strains harboring an expression construct for rituximab in infiltration medium supplemented with kifunensine. After protein purification, N-glycans on the proteins were sequentially modified in vitro by several *E. coli* produced enzymes (depicted in blue boxes in the middle column) for two hours, in either 1:20 (left) or 1:10 (right) enzyme:substrate ratio. Black arrows indicate sequential in vitro reactions. Peaks that could not be labeled to glycans are left unannotated. Glycan structures are drawn according to symbol nomenclature for glycans (SNFG).

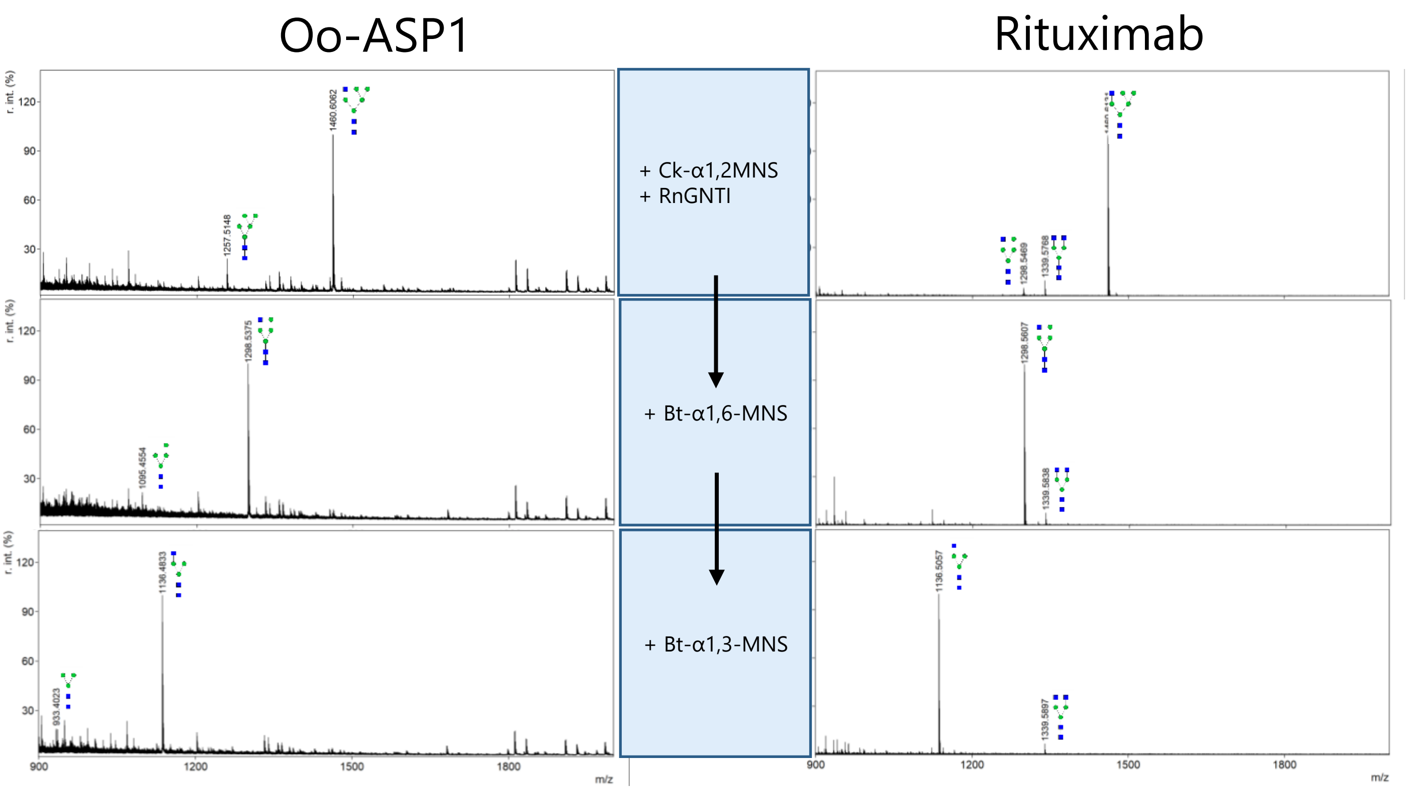


**Supplementary Figure S6: MALDI-TOF MS spectra of GlcNAcMan3GlcNAc2 glycans generated on OoASP-1 and rituximab via the natural glycan processing route.** *N. benthamiana* plants were infiltrated with *A. tumefaciens* strains harboring expression constructs for OoASP-1 (left column) or rituximab (right column) in infiltration medium supplemented with kifunensine. After protein purification, N-glycans on the proteins were sequentially modified in vitro by several *E. coli* produced enzymes (depicted in blue boxes in the middle column). Black arrows indicate sequential in vitro reactions. Peaks that could not be labeled to glycans are left unannotated. Glycan structures are drawn according to symbol nomenclature for glycans (SNFG).

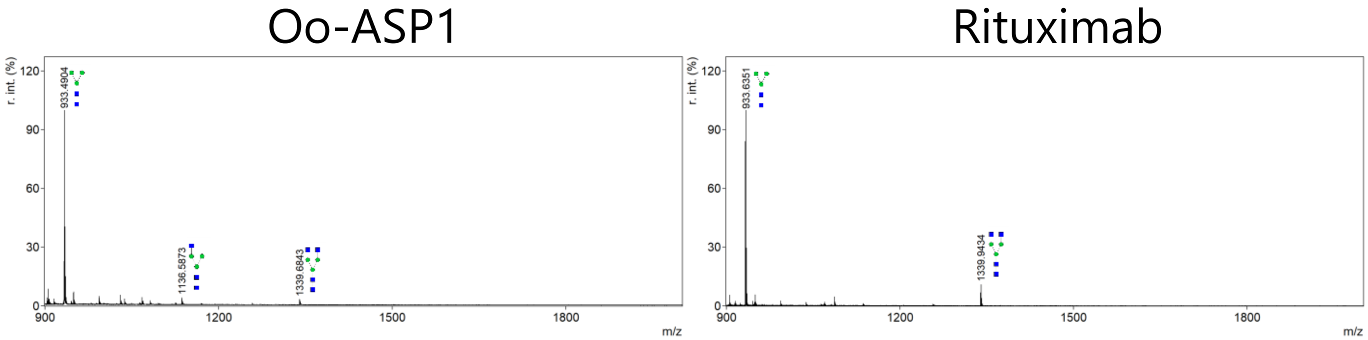


**Supplementary figure S7: MALDI-TOF MS spectra of Man3GlcNAc2 glycans generated on OoASP-1 and rituximab in a one-pot reaction.** *N. benthamiana* plants were infiltrated with *A. tumefaciens* strains harboring expression constructs for rituximab in infiltration medium supplemented with kifunensine. After protein purification, N-glycans on the proteins were modified in a one-pot *in vitro* reaction with *E. coli* produced Ck-α1,2-MNS, Bt-α1,6-MNS and, Bt-α1,3-MNS. Glycan structures are drawn according to symbol nomenclature for glycans (SNFG).

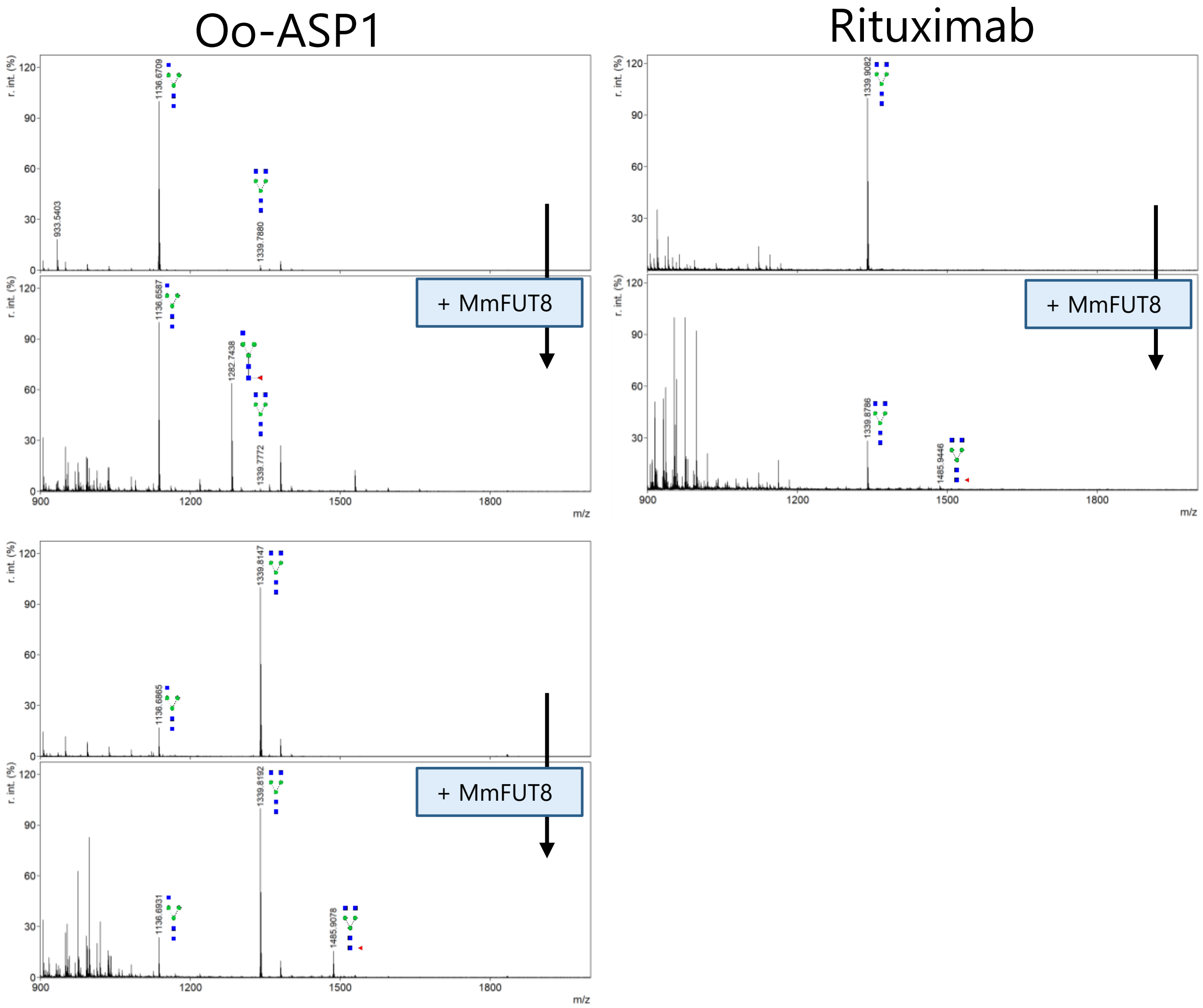


**Supplementary Figure S8: MALDI-TOF MS spectra of *in vitro* fucosylated glycans generated on rituximab.** *N. benthamiana* plants were infiltrated with *A. tumefaciens* strains harboring expression constructs for rituximab in infiltration medium supplemented with kifunensine. After protein purification, N-glycans on the proteins were sequentially modified in vitro by *E. coli* produced MmFUT8. Blue boxes depict enzymes used for in vitro glycoengineering. Black arrows indicate sequential *in vitro* reactions. Peaks that could not be labeled to glycans are left unannotated. Glycan structures are drawn according to symbol nomenclature for glycans (SNFG).

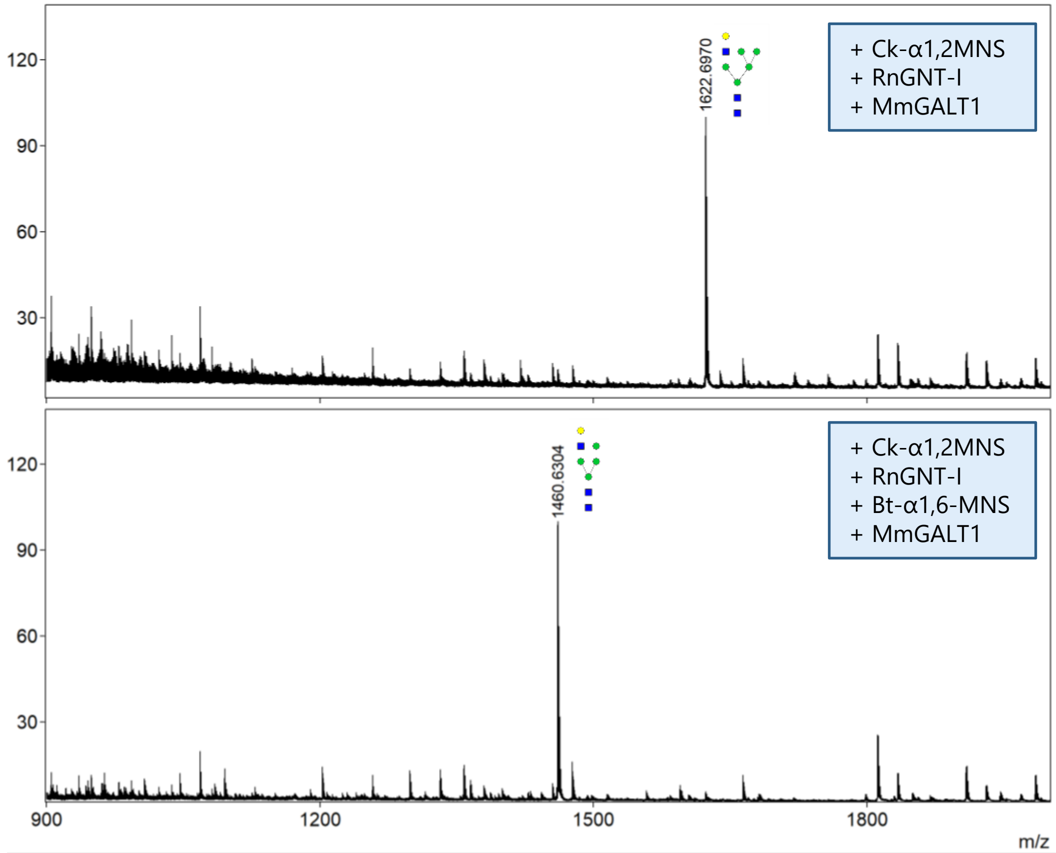


**Supplementary Figure S9: MALDI-TOF MS spectra of unusual galactosylated glycans generated on OoASP-1.** *N. benthamiana* plants were infiltrated with an *A. tumefaciens* strain harboring an expression construct for OoASP-1 in infiltration medium supplemented with kifunensine. After protein purification, N-glycans on OoASP-1 were sequentially modified *in vitro* by *E. coli* produced Ck-α1,2MNS, RnGNT-I and MmGALT1 (**A**) or Ck-α1,2-MNS, Bt-α1,6-MNS, RnGNT-I and MmGALT1 (**B**). Peaks that could not be labeled to glycans are left unannotated. Glycan structures are drawn according to symbol nomenclature for glycans (SNFG).

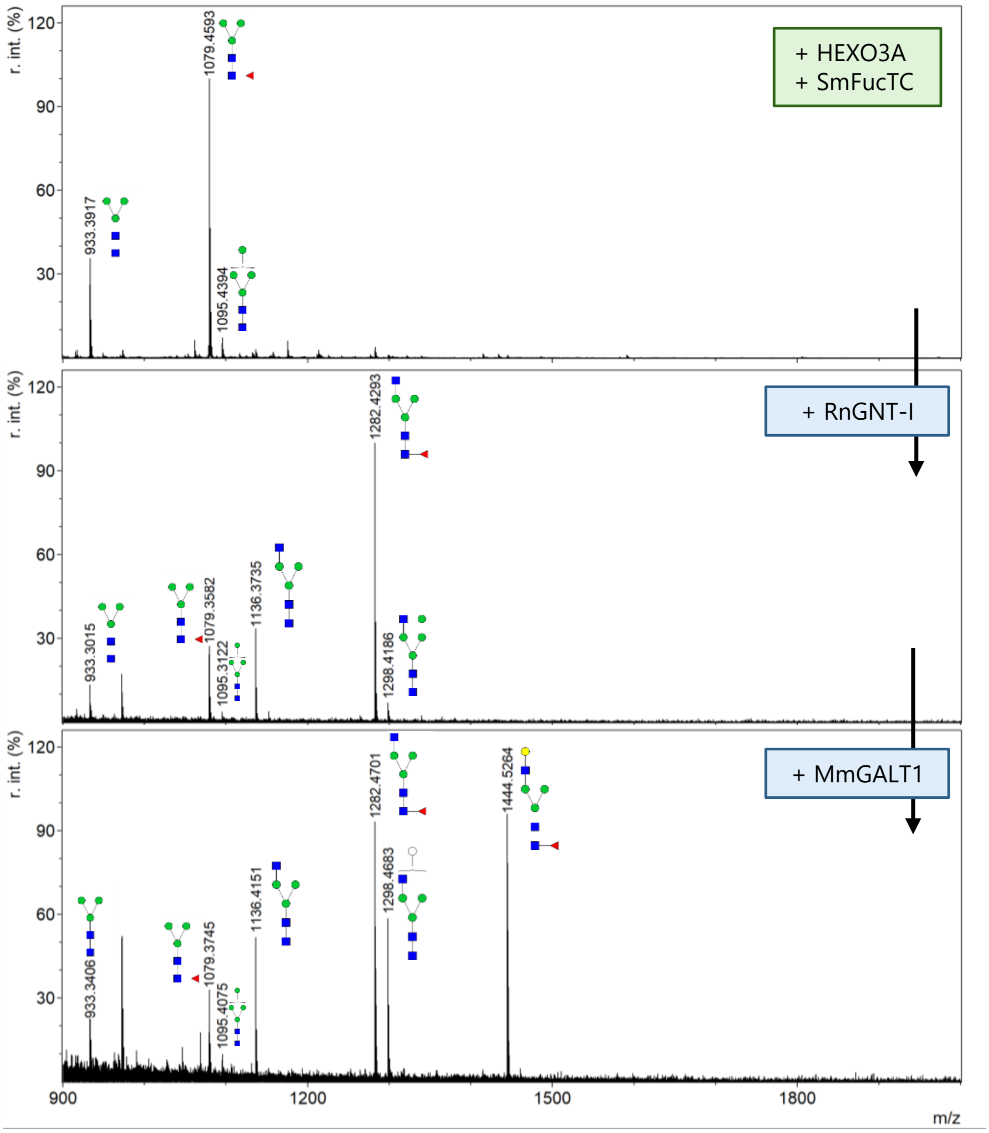


**Supplementary Figure S10. MALDI-TOF MS spectra of PNGase L released glycans generated on OoASP-1 through combining *in planta* and *in vitro* glycoengineering.** *N. benthamiana* plants were infiltrated with *A. tumefaciens* strains harboring expression constructs for OoASP-1, NbHEXO3A and SmFucTC to obtain OoASP-1 with $Man_3GlcNAc_2$ Fuc glycans (**A**). Subsequent *in vitro* glycoengineering with RnGNT-I, RnGNT-II and MmGALT1 was performed to generate native OoASP glycans (**B, C, D, E**). Green boxes depict enzymes used for *in planta* glycoengineering, blue boxes depict enzymes used for *in vitro* glycoengineering. Black arrows indicate sequential *in vitro* reactions. Peaks that could not be labeled to glycans are left unannotated. Glycan structures are drawn according to symbol nomenclature for glycans (SNFG).

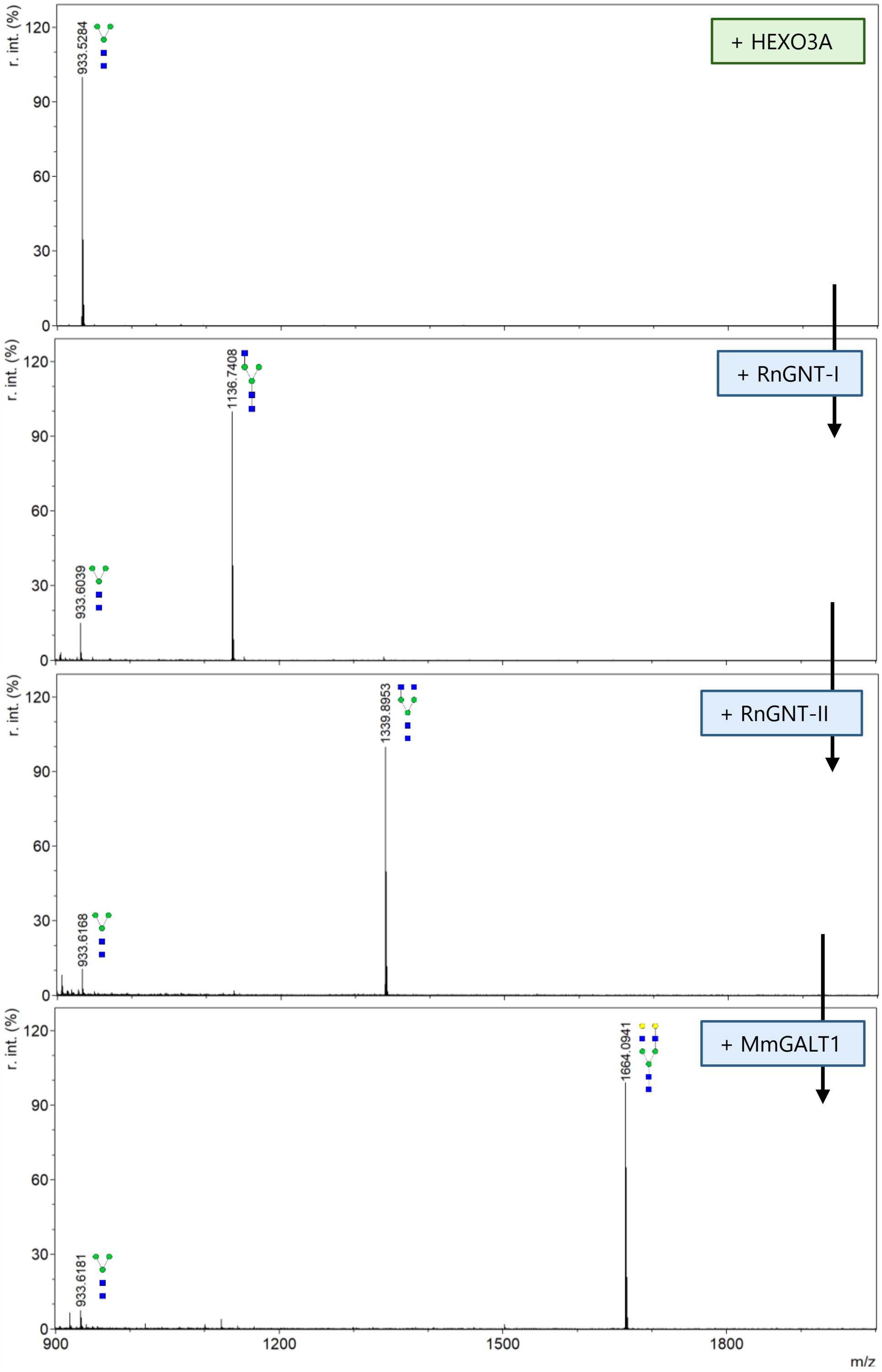


**Supplementary Figure S11. MALDI-TOF MS spectra of PNGase F released glycans generated on OoASP-1 through combining *in planta* and *in vitro* glycoengineering.** *N. benthamiana* plants were infiltrated with *A. tumefaciens* strains harboring expression constructs for OoASP-1 and NbHEXO3A to obtain OoASP-1 with $Man_3GlcNAc_2$ glycans (**A**). Subsequent *in vitro* glycoengineering with *E. coli* produced RnGNT-I, RnGNT-II and MmGALT1 was performed to generate native OoASP glycans (**B, C, D, E**). Green boxes depict enzymes used for *in planta* glycoengineering, blue boxes depict enzymes used for *in vitro* glycoengineering. Black arrows indicate sequential *in vitro* reactions. Peaks that could not be labeled to glycans are left unannotated. Glycan structures are drawn according to symbol nomenclature for glycans (SNFG).
